# Epigenetic Modulation to perturb the *SYNGAP1* Intellectual Disability (ID) that ameliorates synaptic and behavioural deficits

**DOI:** 10.1101/2024.01.03.574003

**Authors:** Akash Kumar Singh, Ila Joshi, Neeharika M. N. Reddy, Sushmitha S. Purushotham, M. Eswaramoorthy, Madavan Vasudevan, Sourav Banerjee, J. P Clement, Tapas K Kundu

## Abstract

Sporadic heterozygous mutations in *SYNGAP1* affects social and emotional behaviour that are often observed in intellectual disability (ID) and autism spectrum disorder (ASD). Although neurophysiological deficits have been extensively studied, the epigenetic landscape of *SYNGAP1* mutation-mediated intellectual disability is unexplored. Here, we have surprisingly found that the p300/CBP specific acetylation marks of histones are significantly repressed in the adolescent hippocampus of *Syngap1^+/-^* mouse. To establish the causal relationship of *Syngap1^+/-^* phenotype and the altered histone acetylation signature we have treated 2-4 months old *Syngap1^+/-^* mouse with glucose-derived carbon nanosphere (CSP) conjugated potent small molecule activator (TTK21) of p300/CBP lysine acetyltransferase (CSP-TTK21). The enhancement of the p300/CBP specific acetylation marks of histones by CSP-TTK21 restored deficits in spine density, synaptic function, and social preferences of *Syngap1^+/-^* mouse that is very closely comparable to wild type littermates. The hippocampal RNA-Seq analysis of the treated mice revealed that the expression of many critical genes related to the ID/ASD reversed due to the treatment of the specific small molecule activator. This study could be the first demonstration of the reversal of autistic behaviour and neural wiring upon the modulation of altered epigenetic modification (s).

## Introduction

Neurodevelopmental disorders (NDD) such as ID and ASD are characterized by poor cognitive, social, emotional, and adaptive functions (*1, 2*). While the precise neurophysiological and neurobiological causes underlying these impairments are understood to an extent, the consensus is that they are a consequence of disrupted neuronal network connections that prevent proper information processing capabilities, particularly during the early stages of development (*3, 4*). The mutations in genes encoding proteins that regulate neuronal connections and processing are increasingly implicated in ID/ASD. Heterozygous mutations in *SYNGAP1*, Synaptic RAS-GTPase activating protein (SYNGAP1), were first reported in 2009 in patients with non-syndromic ID and ASD (*5-7*) and followed by many similar reports (*8-10*). SYNGAP1 is downstream of NMDA receptors and negatively regulates AMPAR insertion via RAS activity (*11, 12*) (*13*), which is crucial for experience dependent synapse function and maturation (*14-17*). Heterozygous mutation in *Syngap1* (*Syngap1^+/-^*) accelerates dendritic spine maturation, altered critical period of plasticity, and sensory processing (*18-21*). Furthermore, the restoration of Syngap1 through genetic means following the critical period of neuronal maturation has been observed to reinstate synaptic function, specifically long-term potentiation (LTP). However, it has not been successful in addressing the associated behavioral deficits (*19, 22-25*). This indicates that the main obstacle lies in the reestablishment of neuronal network connections, which become significantly more resistant to alterations after the critical period of development. (*26*). Therefore, it is crucial to find a therapeutic intervention that may restore neuronal and behavioral functions when the critical period of development is complete.

Recent evidence suggests that the epigenetic processes are dysregulated in neurodevelopmental disorders such as Kabuki syndrome, Rubinstein-Tyabi syndrome RSTS (*27*). These modifications primarily regulate the chromatin dynamics and hence controls the temporal and spatial expression of genes involved in synapse formation, elimination, and maintenance starting from early stages of development. Reversible acetylation of nucleosomal histones, Lysine acetyltransferases (KATs) and Lysine de-acetyltransferases (KDACs), are one of the key epigenetic mechanisms involved in several physiological and pathophysiological processes, including ID/ASD (*28, 29*). The master epigenetic enzymes, p300/CBP were found to be essential KATs for neuronal development as the KAT activity is critical for the differentiation and maturation of neurons (*30*). Moreover, p300 and CBP play important roles in memory and cognitive processes as well (*31-34*). Studies using transgenic mouse models with an inhibitory truncated form of p300 or sub-region specific conditional knock-out of p300 showed deficits in cognition and long-term memory retention (*35*), comparable with Rubinstein-Tyabi syndrome (RTS) (*36-39*). Using zebrafish and mouse model of RTS, studies have shown that restoring acetylation either through genetic rescue of p300/CBP or treatment with KDAC inhibitors such as TSA and SAHA significantly rescued morphological, cognitive, and synaptic plasticity deficits (*40*). Similarly, targeting these modifications instead of synaptic targets are showing promising results in various neurodevelopmental disorders (*27*). However, the epigenetic landscape of *Syngap1* related ID/ASD is not known. Here, we report that there is severe de-regulation of different histone acetylation marks primarily mediated by p300/CBP lysine acetyltransferases (KATs).

While most work to date was focused on targeting synaptic proteins and signalling pathways, little is known about the epigenetic modifications in *Syngap1* and its relationship in causing ID/ASD. We have discovered a specific activator (TTK21) of p300/CBP KATs, which when conjugated to glucose-derived carbon nanospheres (CSP; CSP-TTK21), efficiently delivered to the mice brain, and induced histone acetylation, increased dendritic branching, and extended the retention of spatial memory in normal mice (*41*). CSP-TTK21 has been also used as a therapeutic option in different disease model systems such as in Alzheimer’s Disease and spinal cord injury primarily by restoring the p300/CBP mediated acetylation marks and axonal regeneration respectively (*42*) (*43*). Based on these studies, we administered CSP-TTK21 in adult *Syngap1^+/-^* mice that restored neuronal function and behaviour deficits. The RNA sequence data reveal restoration of gene expression involved in synaptic function and morphology such as Adcy1, FoxJ1, and Ntrk3, but not the reversal of defective *Syngap1*.

## Results

### Activation of p300 KAT activity improves LTP deficits in *Syngap1^+/-^* mouse

While screening the altered epigenetic landscape of *Syngap1^+/-^* mice as compared to the wildtype mice, we observed that *Syngap1^+/-^* mouse possess dramatically low level p300 specific acetylation of histone in the hippocampus (Fig. 1). Both by immunoblotting (Fig. 1, A and B) and immunofluorescence studies (Fig. 1, C and D), we found that histone H3K14 and H4K12 acetylation were significantly reduced, primarily in the hippocampal region of the *Syngap1^+/-^* mice as compared to the control wildtype brain samples. Interestingly, these acetylation levels were associated with transcriptionally active chromatin (*44*), suggesting the possible altered global gene expression in the *Syngap1^+/-^*mouse brain.

**Fig. 1.**
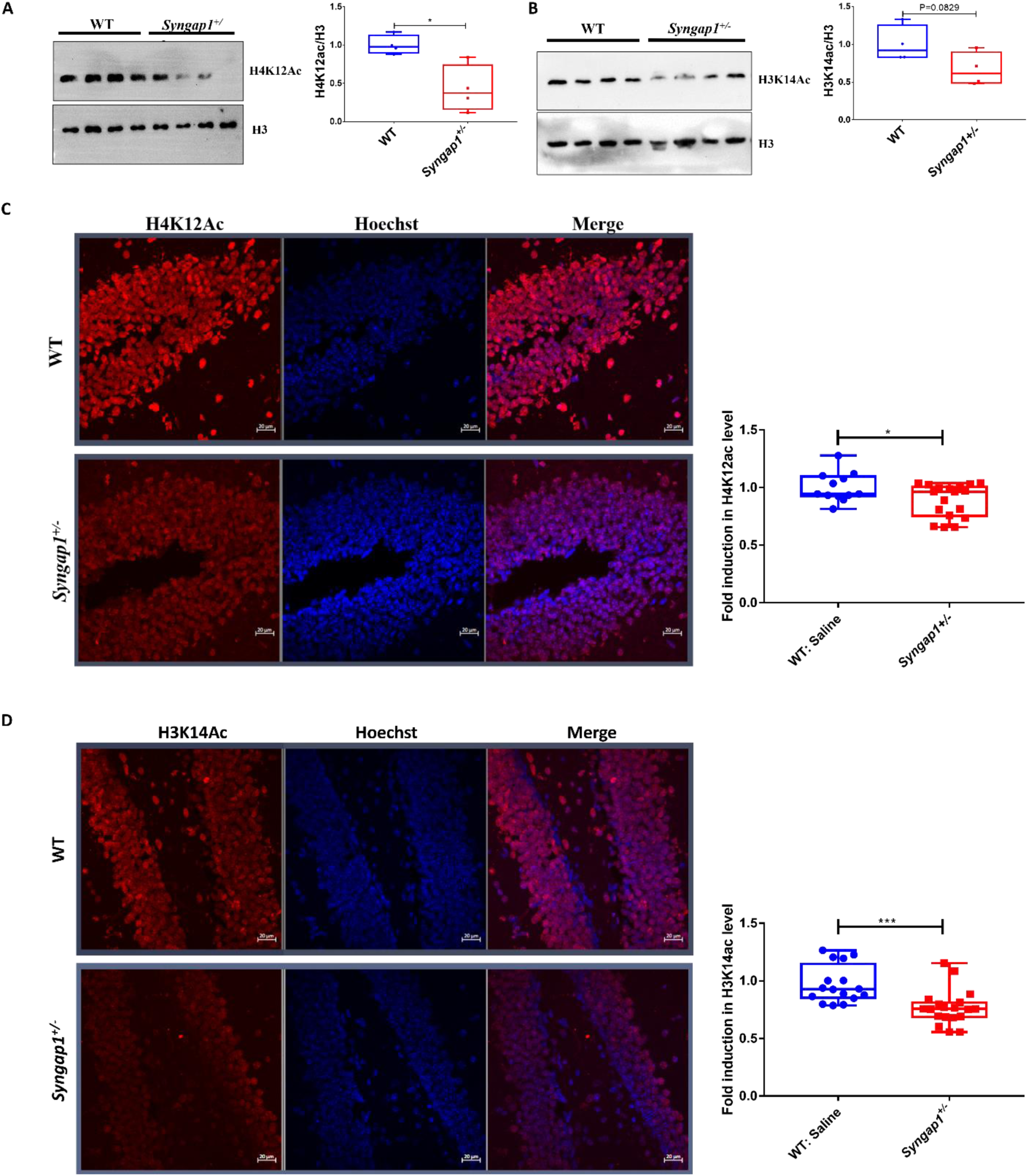
*Syngap1* haploinsufficiency disrupts acetylation machinery. A) Reduced levels of H4K12Ac in *Syngap1^+/-^*mice hippocampus (4 mice/group). B) No change in the levels of H3K14Ac in *Syngap1^+/-^* mice hippocampus (4 mice/group). C) Representative fluorescence image showing H4K12Ac level in dorsal hippocampus, depicting reduced levels in *Syngap1^+/-^* mice (WT, n=12 sections, 4 mice; *Syngap1^+/-^*, n=18 sections, 6 mice; unpaired *t-test*). D) Reduced H3K14Ac levels in the dorsal hippocampus of *Syngap1^+/-^* mice as compared to WT mice, representing disruption of acetylation machinery (WT, n=16 sections, 4 mice; *Syngap1^+/-^*, n=19 sections, 6 mice; unpaired *t-test*). Scale 20 µm, error bars represents the standard error of mean (s.e.m). *p<0.05, **p<0.01 and ***p<0.001.

The reduced p300 KAT activity in *Syngap1^+/-^* mouse brain, encouraged us to investigate the efficacy of small molecule activator of p300 to rescue the neurophysiological deficits associated with *Syngap1^+/-^*mouse model, primarily through the restoration of acetylation machinery.

In earlier studies we have found a specific p300/CBP KAT activator, TTK21, conjugated to glucose derived carbon nanosphere (CSP), CSP-TTK21 could induce histone acetylation and adult hippocampal neurogenesis in adult mice (*41*). We have found that CSP-TTK21 could be delivered orally and is as effective as IP administration (Fig. S2) (*45*). Thus, in this study, we conducted an oral gavage treatment on five distinct groups of adult mice. Each group received one dosage of 20 mg/kg body weight of CSP-TTK21 and tested 84 hours after the treatment (Fig. 2A). The groups were as follows: wild type (WT) mice treated with saline (WT), *Syngap1^+/-^* mice with saline (*Syngap1^+/-^*), *Syngap1^+/-^* mice with CSP-vehicle control (*Syngap1^+/-^*: CSP), *Syngap1^+/-^* with CSP- TTK21 (*Syngap1^+/-^*: CSP-TTK21), and WT with CSP-TTK21 (WT: CSP-TTK21) for 84 hrs. The animals were then sacrificed and subjected to immunohistochemistry analysis. We observed a significant increase in H4K12Ac and H3K14 Ac levels in CSP-TTK21 treated group as compared to CSP and saline-treated groups (Fig. 2B and S3). Significantly, the acetylation levels were nearly restored to WT group upon the treatment with CSP-TTK21 (Fig. 2B and S3). Since p300/CBP has been linked with neuronal proliferation and maturation (*30*), and *Syngap1^+/-^* mice show accelerated critical period development (*19*), we next asked whether the generation of new born neurons are altered in these mice. We observed a relatively a smaller number of DCX-positive immature neurons primarily in the dentate gyrus region of the hippocampus (Fig. 2C). However, CSP-TTK21 treatment led to an increase in the number of DCX^+^ immature neurons as compared to CSP and saline control groups (Fig. 2C).

**Fig. 2.**
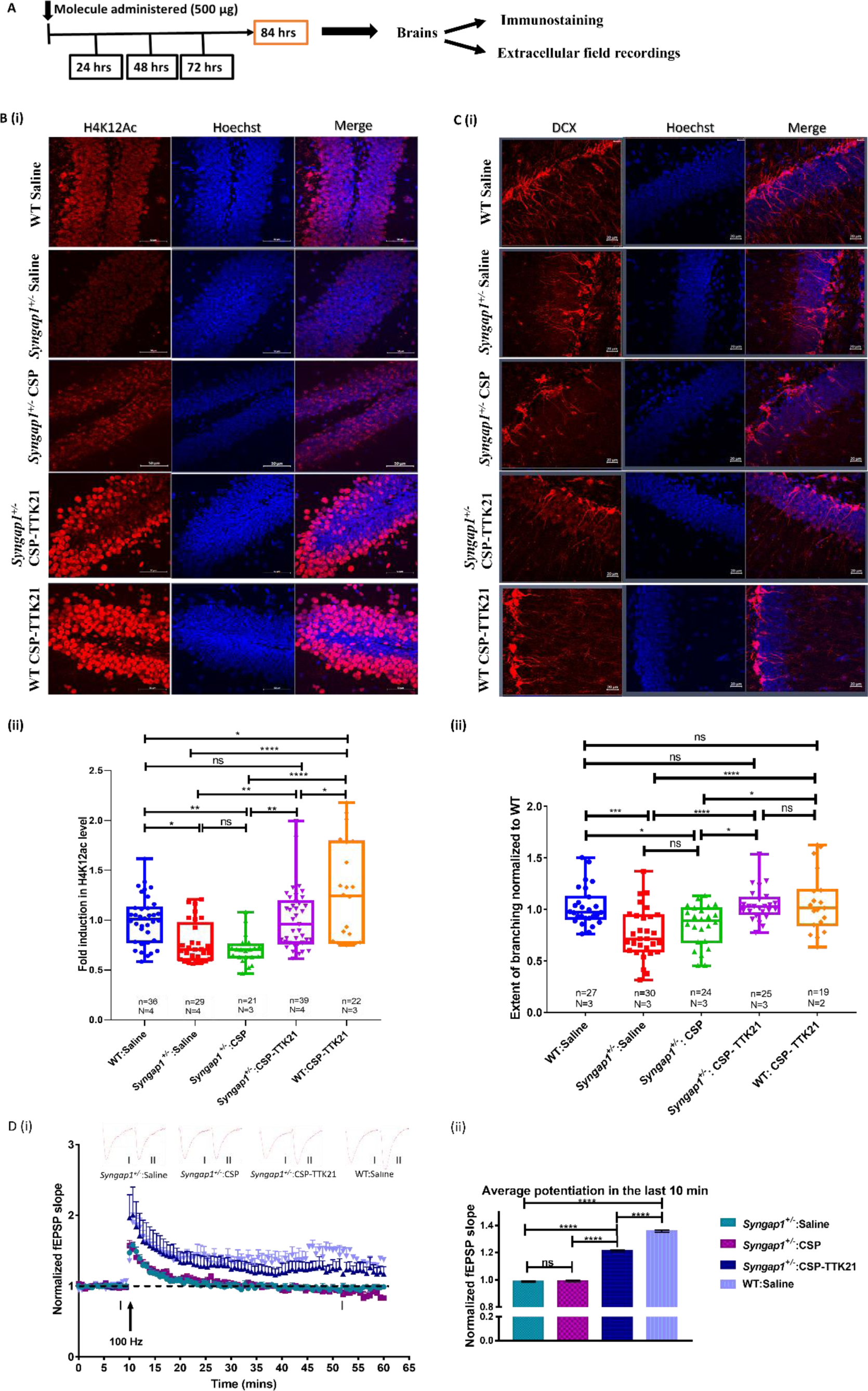
Activation of p300/CBP KAT activity restores synaptic plasticity through restoration of acetylation levels and increased dendritic branching. A) Experimental schematic, mice were sacrificed after 84 hours of treatment and experiments were performed. B) Representative confocal images showing H4K12Ac expression in dorsal hippocampus of the different experimental groups used (*Syngap1^+/-^*:Saline, *Syngap1^+/-^*:CSP, *Syngap1^+/-^*:CSP-TTK21, and WT:CSP-TTK21) depicting reduced levels in *Syngap1^+/-^* mice treated with saline or CSP, and restoration to WT levels in CSP-TTK21 treated *Syngap1^+/-^*mice. CSP-TTK21 treated WT mice also showed increased levels of H4K12Ac as compared to control WT, depicting CSP-TTK21 functional activity. C) Representative confocal images of DCX^+^-neurons in the subventricular zone of Hippocampus in the treatment groups, depicting reduced dendritic braching in saline or CSP treated *Syngap1^+/-^* mice, and restoration to WT levels in CSP-TTK21 treated *Syngap1^+/-^* mice. Scale 20 µm, error bars represent the s.e.m. D) Extracellular field recoding from acute hippocampus slices, showing impaired LTP induction in *Syngap1^+/-^*mice treated with saline or CSP. CSP-TTK21 improves LTP induction, didn’t return to baseline till the end of experiment (60 min). Example traces are average of those recorded in 1–2 min around the time point indicted (I and II). Qunatification showing significant changes in the last 10 min of recording (WT, n=4 slices, 3 mice; all other groups n=6 sections, 3 mice each). One-way ANOVA with Turkey’s post-hoc test was perfomed for statistical analysis, *p<0.05, **p<0.01 and ***p<0.001.

Recent evidence suggests that these immature neurons are crucial for learning and memory processes (*46*), and can integrate into the experience dependent functional neurons in the existing or a novel neuronal networks (*47, 48*).

Taken together, these results suggest that CSP-TTK21 by activating p300/CBP KAT activity could increase DCX^+^ immature neurons in *Syngap1^+/-^*mice primarily through the restoration of acetylation marks.

Earlier studies report altered neuronal network and deficits in synaptic plasticity in *Syngap1^+/-^*mice (*18-21*). We further asked the question whether these newly formed neurons can integrate in the neuronal network in response to activity, and thereby restore the synaptic plasticity. Therefore, we next investigated the effect of CSP-TTK21 on the neuronal network and synaptic plasticity in the hippocampal region of these mice. For this purpose, we performed extracellular field recordings in the Schaffer-Collateral pathway from acute brain hippocampal slices.

As expected, we observed that the long-term potentiation (LTP) in CSP-TTK21 treated *Syngap1^+/-^* as compared to saline/CSP treated group that lasted until the end of recordings (1 hr) and was restored to the WT level (Fig. 2D and S4). These results further suggest that CSP-TTK21 could rescue synaptic deficits associated with *Syngap1^+/-^*mice by increasing the number of newborn neurons that subsequently incorporated in the functional neuronal networks.

### Does p300 KAT activity induced recovery of long-term potentiation (LTP) deficit affects the *Syngap1^+/-^* mice behaviour?

Previous studies reported that the LTP in adult *Syngap1^+/-^* mice were restored to the WT levels but did not rescue various behavioural deficits in *Syngap1^+/-^* mice (*19, 22-25*). Encouraged by the LTP restoration upon the treatment of the p300 KAT activator, we asked if p300/CBP KAT activation by CSP-TTK21 rescued behavioural deficits observed in *Syngap1^+/-^* mice. For this purpose, we performed various behavioural paradigm studies described in Fig. 3a. Briefly, mice were grouped as described earlier and treated for 3 weeks (1 dosage/week) with Saline or CSP-TTK21 before behavioural assays. WT littermates treated with saline were used as controls.

**Fig. 3.**
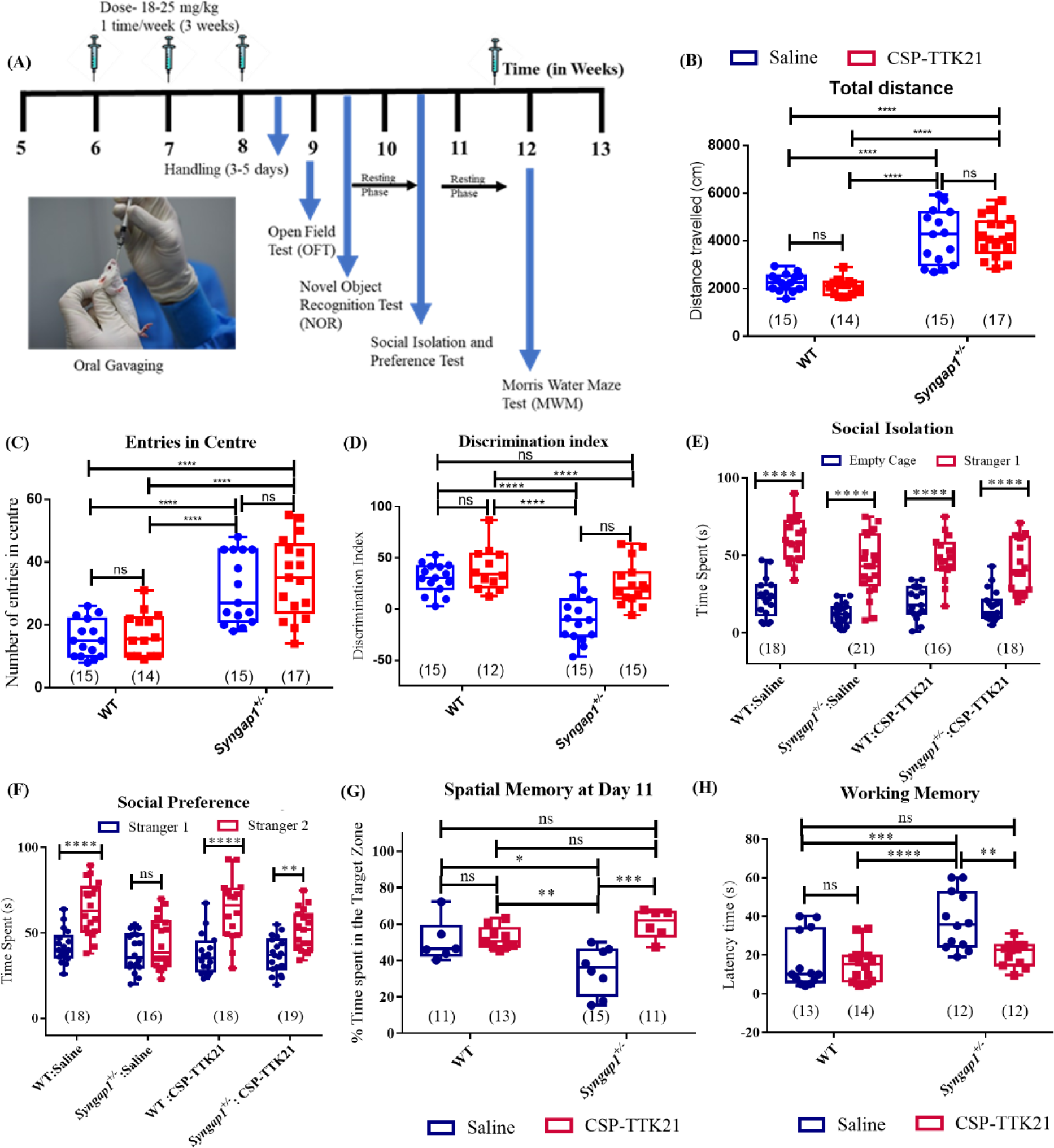
Increasing p300/CBP activity improves behavioural measures. A) Schematic showing experimental strategy. B) Total distance covered in OFT, depicting hyperactivity in *Syngap1^+/-^*mice and no effect of CSP-TTK21 tretament. C) Number of entries in centre in OFT, depicting anxiolytic behaviour in *Syngap1^+/-^* mice and no effect of CSP-TTK21 tretament. D) Spatial memory at Day11 of training (72 hours after the last training session), *Syngap1^+/-^* mice show reduced % time spent in the target zone indicating impaired spatial memory. CSP-TTK21 treatment led to increase % time spent in the target zone implicating improved spatial memory. E) Latency time for working memory, *Syngap1^+/-^*mice show increased latency time depicting their inability to learn the new location of the platform which was changed daily. CSP-TTK21 treatment led to decrease latency time in *Syngap1^+/-^* mice. F) Time spent with empty cage or stranger mice, *Syngap1^+/-^*mice as well as WT mice shows increased interaction with stranger1 mice indicating no deficits in social interaction. G) Time spent with stranger1 or stranger2 (novel) mice, *Syngap1^+/-^*mice show reduced interaction with stranger2 mice as compared to WT mice indicating deficits in social preference. CSP-TTK21 treatment led to increased interaction of *Syngap1^+/-^* mice with stranger2 mice. Error bars represent s.e.m. Two-way ANOVA with bonferonni’s post-hoc was done for significance test, *p<0.05, **p<0.01 and ***p<0.001.

It is known from earlier studies that *Syngap1^+/-^*mice are hyperactive and less anxious in an open field arena (*17, 49*). Our results from open field arena test concurs with the previous published studies where *Syngap1^+/-^* mice travels significantly longer distance and showed increased entry in the centre of the arena as compared to WT controls (Fig. 3, B and C, and S5A). However, treatment of CSP-TTK21 did not rescue the locomotor and anxiety related issue of *Syngap1^+/-^* mice. The CSP-TTK21 did not affect the WT mice (Fig. 3, B and C).

It is shown that the *Syngap1^+/-^*mice are unable to distinguish between the familiar and the new objects, thereby, alluding impaired memory (*17, 49*). Similarly, we also observed that *Syngap1^+/-^* mice were unable to discriminate between a familiar object and a novel object after 24 hours of familiarisation as compared to WT mice (Fig. 3D). Significantly, upon the CSP-TTK21 treatment, *Syngap1^+/-^* mice were able to efficiently discriminate between the two objects, as shown by the discrimination index (DI) (Fig. 3D). The recognition index (RI), a measure of recognition memory and the DI of CSP-TTK21 treated *Syngap1^+/-^* mice group were found to be similar to the WT group (Fig. S5B and Fig. 3D), indicating significant improvement and restoration of this phenotype.

One of the major behaviour dysfunctions observed in children with autism is lack of sociability (*50, 51*). Further, we wanted to investigate the social behaviour in *Syngap1^+/-^*mice. For this, we used a 3-chamber assay, classic test to assess social behavior and memory in rodents. The aim of this test is to study the sociability and preference for social novelty compared to an object or a familiar mouse, and therefore social memory. Studies from other laboratories have shown that *Syngap1^+/-^* mice display deficits in sociability and prefers not to interact with a novel mouse (*17, 49*). In our experiment, we observed that both the *Syngap1^+/-^* mice and WT mice spent more time with stranger mice cage as compared to empty cage (Fig. 3E). However, when we assessed social preference – where stranger 1 is a familiar mouse and stranger 2 is novel mouse, we observed that *Syngap1^+/-^* mice showed severe impairment in interaction with stranger 2 (novel) mouse than stranger 1(familiar) mouse, as compared to WT mice (Fig. 3F). This lack of social preference was restored in the CSP-TTK21 treated *Syngap1^+/-^* mice to that of saline treated mice (Fig. 3F). Next, we wanted to check the hippocampus-dependent learning and working memory in these mice. For that, we used **Morris Water Maze (MWM)**, a classical test to measure spatial reference and working memory. Mice were trained for 8 days in the presence of distal extra-maze cues to locate a hidden platform. During spatial training, a highly significant decrease in escape latencies (time taken to reach the platform) across days was observed in WT mice as compared to *Syngap1^+/-^* mice which showed a decrease in escape latency. We observed that *Syngap1^+/-^*mice treated with CSP-TTK21 showed decrease in latency similar to that of WT mice, in contrast to the Saline treated *Syngap1^+/-^* mice (Fig. S5C). After 24 hr, a probe test was performed to test spatial reference memory. We observed that *Syngap1^+/-^* mice were not able to remember the target quadrant, where the escape platform was kept during spatial training, as depicted by less time spent in the target quadrant, compared to the WT mice. The CSP-TTK21 group showed increased exploration and time spent in the target quadrant similar to WT mice (Fig. 3G and S5D), suggesting restoration of spatial reference memory. To evaluate the working memory of *Syngap1^+/-^* mice, we subjected these mice to probe test 24-hours after exposing them only once to remember the location of escape platform prior to shifting it to a new location. Testing was performed after 30s of training to analyse the escape latencies which correspond to working memory.

We found severe impairment of working memory in *Syngap1^+/-^* mice as these mice took a long time to reach the platform as compared to the WT mice. However, CSP-TTK21 treated group showed a significant decrease in escape latencies as compared to *Syngap1^+/-^* mice, and the decrease was similar to the WT group (Fig. 3H). Therefore, these results suggest that small activator of p300/CBP, could be a potential therapeutic option in *Syngap1* and other related IDs.

### CSP-TTK21 may lead to rewiring of neuronal networks in the thalamocortical region

Previously, it has been reported that, upon whisker deprivation during critical developmental period, *Syngap1^+/-^* mice had fewer filopodia in the somatosensory cortex as compared to WT animals (*20*). This suggests that, unlike WT mice, *Syngap1* mutation prevents neurons from being plastic and adapting to changes in the sensory input, and once the hard-wiring of connections are established, it is extremely difficult to induce rewiring in the *Syngap1^+/-^* mice.

The reversal of LTP and behavioural deficits by the small molecule activator, TTK21 (conjugated to carbon nanosphere) encouraged us to investigate the functional mechanisms of the reversals these physiological deficits. We investigated whether the activator is inducing the rewiring of the neuronal network that enables them to perform better in the behavioural paradigms tested.

To investigate the functional re-wiring of synaptic connections in the somatosensory cortex, we performed whisker deprivation from PND7-21 and treated these mice with Saline, CSP and CSP- TTK21 as shown in (Fig. 4A). In agreement with previous reports (*20*), we observed a signification increase in filopodia density in trimmed WT mice (WT: Saline, trimmed) as compared to untrimmed WT group (WT: Saline, untrimmed) (Fig. 4, B and D). Furthermore, whisker trimming had no effect on *Syngap1^+/-^* mice (Saline and CSP) as compared to untrimmed *Syngap1^+/-^* mice (Fig. 4D). Interestingly, *Syngap1^+/-^* mice treated with CSPTTK21 showed increased filopodia density similar to that of trimmed WT mice (Fig. 4, B and D). We did not observe any significant effect of whisker trimming in total spine density across various treatment groups (Fig. 4C). Collectively, these encouraging results implicate that CSP-TTK21 might be able to promote synaptic rewiring when administered after the critical period of development and enables the capability to reorganise cortical circuits in response to change in the sensory stimuli.

**Fig. 4.**
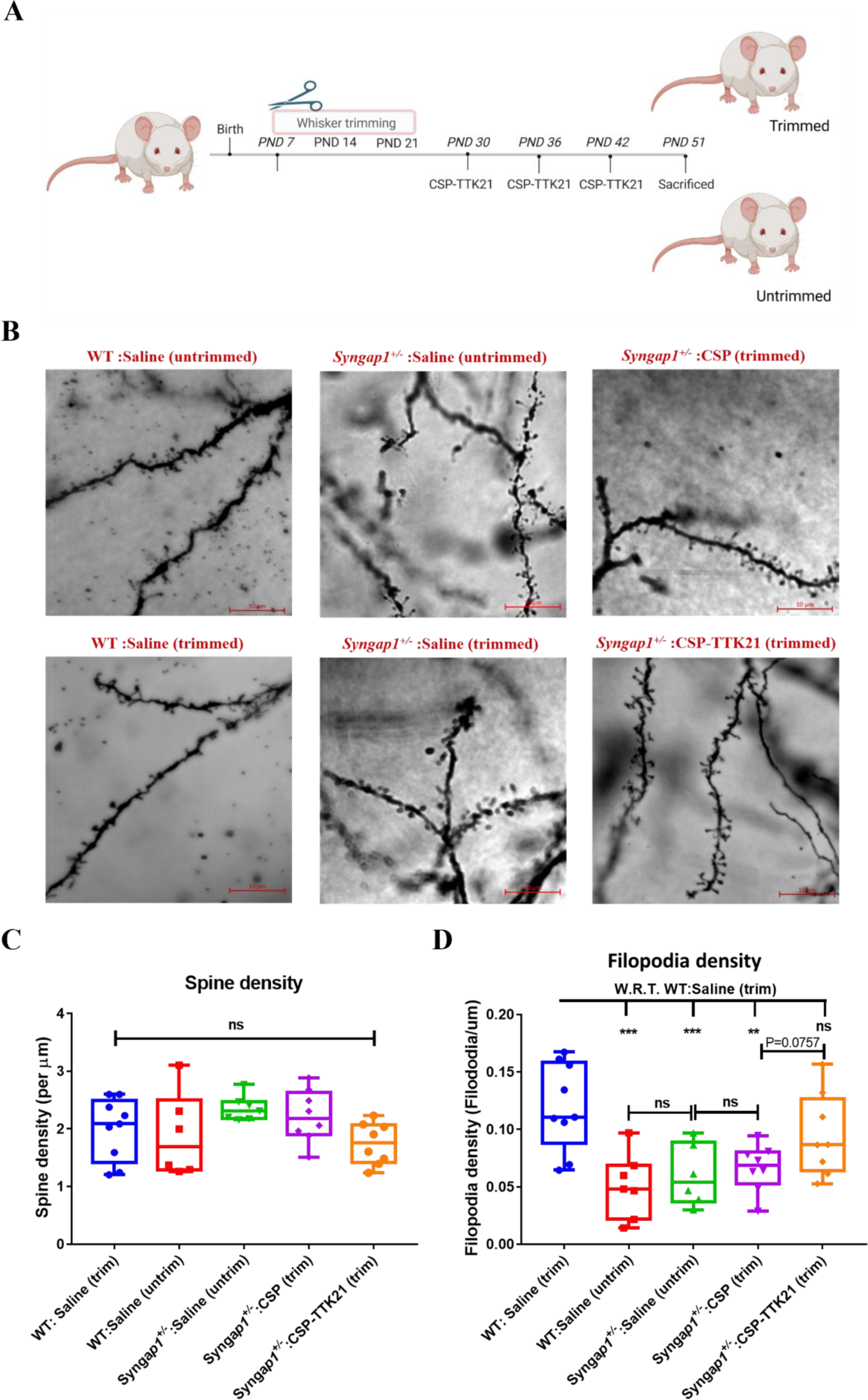
CSP-TTK21 improves synaptic networks in somatosensory cortex: (A) Schematic showing the experimental timeline: whisker trimming was done daily from PND7-PND21, and three dosage of CSP-TTK21 was administered weekly starting from PND30. 3 days after the 3rd dosage mice were sacrificed and Golgi-staining was performed. (B) Representative images from each group. Scale 10μm. (C) Quantifications for spine density and (D) filopodia density across groups (n=6-12 sections and N= 2/3 mice). One-way ANOVA with Turkey’s post-hoc test was perfomed for statistical analysis.

### 5. Molecular pathways involved in the p300 activation mediated rescue of several neurological deficits in *Syngap1^+/-^* mice

To understand the underlying molecular mechanisms in the rescue of ID in *Syngap1^+/-^* mice, we investigated the direct effect of the activator, on SYNGAP1 and its downstream pathway. For this purpose, we checked the expression of SYNGAP1 and its downstream target p-ERK/ERK by western-blotting from Hippocampal lysates upon the treatment of CSP-TTK21. Interestingly, we did not find any significant difference in the levels of either of the proteins in *Syngap1^+/-^*mice treated with saline, CSP or CSP-TTK21 (Fig. S6). These results demonstrate that CSP-TTK21 is not directly acting on the SYNGAP1 or its related pathway, but through some other complimentary network to rescue the deficit. Therefore, to delineate the molecular networks through which this small molecule activator could be acting, we performed whole hippocampal transcriptome analysis across 6 treatment groups (1. WT: Saline, 2. *Syngap1^+/-^*: Saline, 3. WT: CSP, 4. *Syngap1^+/-^*: CSP, 5. WT: CSP-TTK21 and 6. *Syngap1^+/-^*: CSP-TTK21).

Interestingly, we observe 2600 differentially expressed genes (DEGs) in *Syngap1^+/-^* mice with respect to WT mice (*Syngap1^+/-^*: Saline Vs WT: Saline) of which 2098 genes were down regulated, and 502 genes were upregulated (Fig. 5Ai). Upon GO and pathway annotation we found majority of DEGs are involved in mRNA processing, signalling processes including WNT, AKT, ErbB, VEGF, NOTCH, etc., and various synaptic processes such as synaptic vesicle cycling, neurotransmitter release and voltage-gated ion-channel activity (Fig. 5Bi). Furthermore, when we compared the transcriptomes of *Syngap1^+/-^*: CSP-TTK21 mice (treatment group) with *Syngap1^+/-^*: Saline animals (pathological group), we found 1934 DEGs. Of the 1934 DEGs, 471 genes showed down-regulation and 1463 genes showed up-regulation, demonstrating the positive effect of CSP- TTK21 treatment (Fig. 5Aii). Likewise, when we compared *Syngap1^+/-^*: CSP-TTK21 mice to the vehicle control *Syngap1^+/-^*: CSP mice, we found 2269 DEGs. Of 2269 DEGs, 637 were down-regulated and 1622 were up-regulated, showing the direct effect of CSP-TTK21 as opposed to the CSP vehicle treatment (Fig. 5Aiii). Furthermore, upon GO and molecular pathways annotation, we observed majority of DEGs include genes involved in mRNA processing, synaptic vesicle trafficking, neurotransmitter release, axonal transport, dendritic morphogenesis, chromatin organization and various signalling pathways including WNT, AKT, VEGF, ErbB, etc. which was similar to the *Syngap1^+/-^*: Saline Vs WT: Saline (Fig. 5bii and S8). When we compared the DEGs in the pathological condition (*Syngap1^+/-^*: Saline Vs WT: Saline) with DEGs in treatment condition (*Syngap1^+/-^*: CSP-TTK21 Vs *Syngap1^+/-^*:Saline) we observed that CSP-TTK21 treatment leads to restoration of several DEGs. Specifically, it rescued (increased) expression of 696 genes out of 2098 downregulated genes and decreased expression of 140 genes out of 502 Up-regulated genes (Fig. S7).

**Fig. 5.**
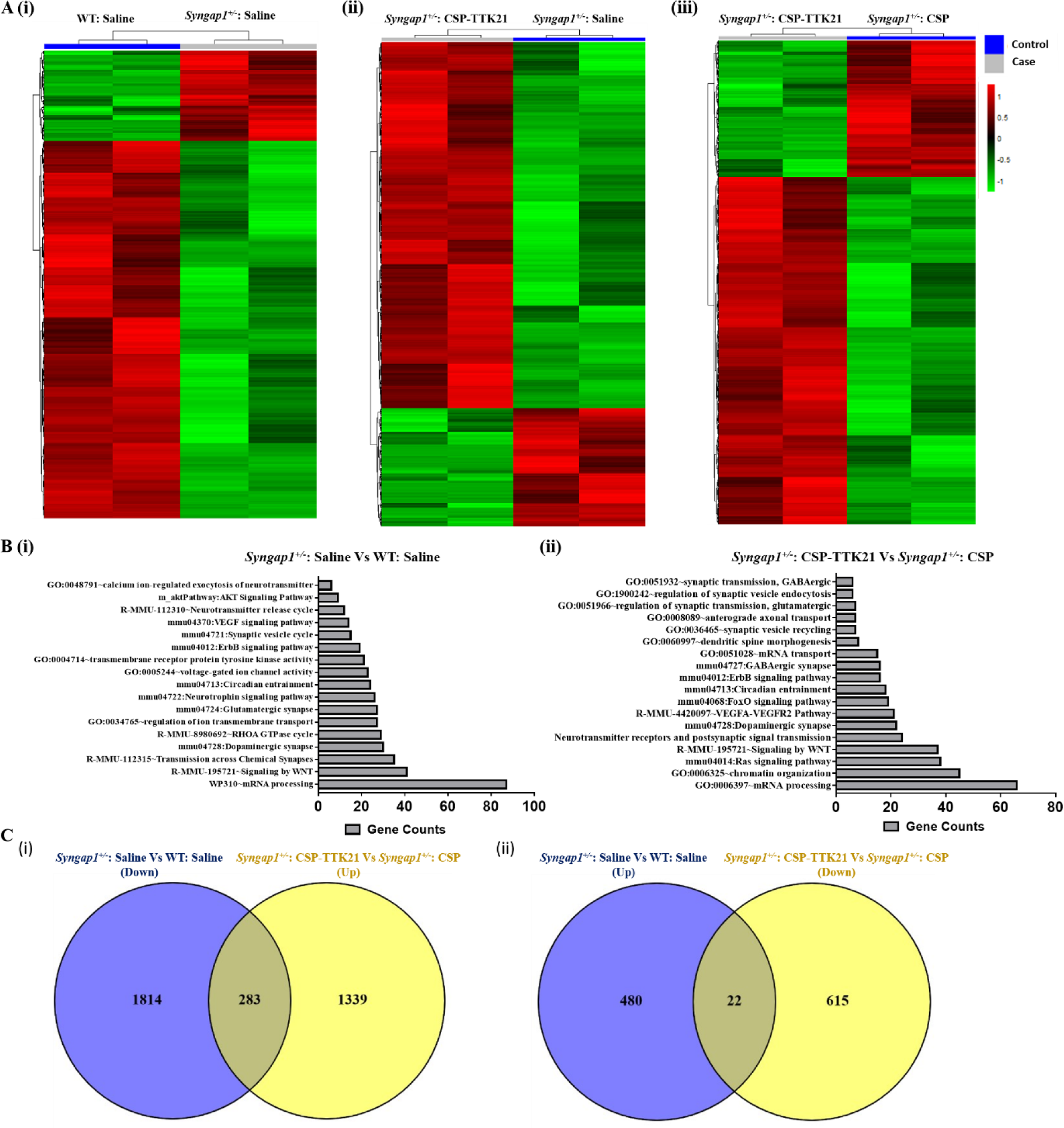
Transcriptomic alteration in *Syngap1^+/-^* mice and effects of CSP-TTK21. A. Heatmaps representing the differentially expressed transcripts across various conditions. (i) *Syngap1^+/-^*: Saline Vs WT: Saline, (ii) *Syngap1^+/-^*: CSP-TTK21 Vs *Syngap1^+/-^*: Saline, and (ii) *Syngap1^+/-^*: CSP-TTK21 Vs *Syngap1^+/-^*: CSP. Blue colour represents the control group and grey represent the case group (effect). Green represents down-regulation and red represents up-regulation. B) GO and molecular pathways annotation of highly significant DEGs. (i) *Syngap1^+/-^*: Saline Vs WT: Saline and (ii) *Syngap1^+/-^*: CSP-TTK21 Vs *Syngap1^+/-^*: Saline. C) Venn diagram showing effect of CSP-TTK21 in restoration of significantly deregulated genes, (i) down-regulated in *Syngap1^+/-^*mice (blue) and up-regulated in *Syngap1^+/-^*: CSP-TTK21 mice (yellow)**. (ii)** Up-regulated in *Syngap1^+/-^* mice (blue) and down-regulated in *Syngap1^+/-^*: CSP-TTK21 mice (yellow). The overlap region (yellowish blue) depicts the restoration of DEGs upon CSP-TTK21 treatment.

Similarly, when we compared the DEGs in pathological condition (*Syngap1^+/-^*: Saline Vs WT: Saline) with DEGs in the other treatment condition (*Syngap1^+/-^*: CSP-TTK21 Vs *Syngap1^+/-^*: CSP), we observed restoration of several DEGs upon CSP-TTK21 treatment. Specifically, it rescued (increased) expression of 283 gene out of 2098 down-regulated genes and reduced expression of 22 genes out of 502 Up-regulated genes (Fig. 5C). Furthermore, upon molecular network analysis we observed inverse correlation for most of the affected molecular pathways in these conditions. Genes that were upregulated or downregulated in pathological conditions their expression patterns were reversed upon CSP-TTK21 treatment (Fig. 6, A and B). Besides, we found several important neuronal genes crucial for diverse biological processes including synaptic plasticity, learning and memory, dendritic spine morphogenesis and axonal transport which were deregulated in the *Syngap1^+/-^* mouse were rescued by CSP-TTK21 treatment. Furthermore, we verified these findings by using Real-time Polymerase Chain Reaction (RT-PCR) to measure the expression of DEGs. As compared to WT group, we observed down-regulation of Adcy1, Egr1, Erbb3, Foxj1, Gng7, Kcnq3, Kalrn1, Ntrk3 and up-regulation of Gfap, Itpr1, and Npy1 in *Syngap1^+/-^* mice either treated with saline and CSP. Interestingly, CSP-TTK21 treatment restored expression of most of these genes to WT levels (Fig. 6C).

**Fig. 6.**
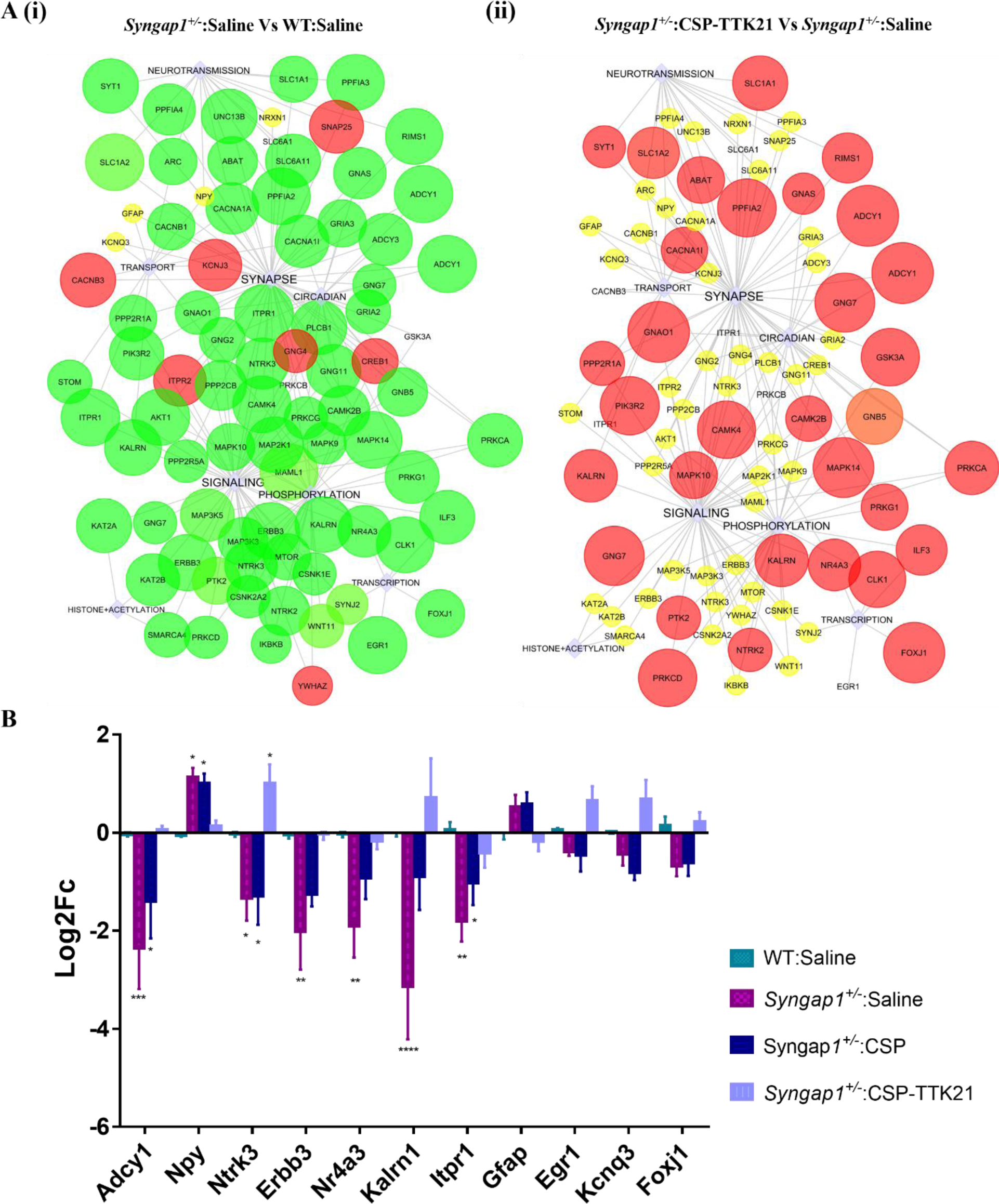
CSP-TTK21 restores gene expression and signalling pathways that were altered in *Syngap1^+/-^* mouse. (A) Significantly de-regulated pathways *Syngap1^+/-^* mouse (i) *Syngap1^+/-^*:Saline Vs WT: Saline. (ii) Significantly restored pathways upon CSP-TTK21 treatment (*Syngap1^+/-^*:CSP-TTK21 Vs *Syngap1^+/-^*: Saline). The circles indicate differentially expressed genes, and boxes indicate pathways regulated by the DEGs. Up-regulated genes are colored in red and down-regulated are in green. Size of the circle indicates p-value (the bigger the size, the lower the p-value). (B) RT-PCR validation of some of the crucial neuronal genes across the different treatment groups (n = 5-6 mice/group). Two-way ANOVA with correction for multiple comparison test was performed for statistical analysis, *p ˂ 0.05, **p ˂ 0.01, and ***p ˂ 0.001.

These observations suggests that CSP-TTK21 restores transcriptomic changes in *Syngap1^+/-^*mice by p300/CBP- mediated activation of histone acetylation marks.

### Discussions and future perspectives

One of the most important findings of the present study is the dramatic reduction of acetylation marks primarily mediated by p300/CBP KATs in the brain of *Syngap1^+/-^*mice.

p300/CBP has been known to play a crucial role in maturation and proliferation of neurons during brain development (*30, 52*). We have also shown that there is a reduced number of DCX^+^ immature neurons in the hippocampal region of these mice. These findings suggest that p300/CBP mediated acetylation might be another crucial factor in bringing upon the associated plasticity and behavioural deficits seen at adulthood and could be a potential therapeutic target. We, therefore, were highly encouraged to test the effect of a specific activator of p300/CBP, TTK21, originally was discovered as CTPB (*53*).

Remarkably, we observed that CSP-TTK21 can restore the acetylation levels as well as DCX^+^ immature neurons in *Syngap1^+/-^* mice to WT levels. Furthermore, we demonstrated that CSP- TTK21 not only rescues the synaptic plasticity deficits but also rescued the social and behavioural abnormalities upon the administration of the activator, after the critical period of development, when the neuronal networks are hard-wired. We also uncovered some of the molecular and physiological changes underlying the rescue effect.

Transcriptomic analysis revealed significant alteration in gene expression in *Syngap1^+/-^* mice as compared to WT mice. This was expected as deficits in synaptic plasticity, spine density and behavioural measures are already well associated with altered gene expression (*54-56*). Our functional enrichment analysis further provide evidence that these altered genes code proteins that play important role in ion transport, neurotransmission, cellular signalling, transcription regulation, and synaptic function. Presumably, these altered genes are the primary source of behavioural abnormalities and synaptic dysfunction in *Syngap1^+/-^* mice. Importantly, CSP-TTK21 treatment partially rescued expression of these genes to WT levels. The CSP-TTK21-mediated significant restoration of Adcy1, Ntrk3, Egr1 and Foxj1 in particular, could have contributed to the functional rescue of LTP, synaptic functions, thalamocortical rewiring and dendritic branching leading to partial restoration of behavioural phenotype in *Syngap1^+/-^* mice. Adcy1 is required for the induction and maintenance of LTP in the hippocampus as well as thalamocortical synapse maturation and aspects of layer IV barrel development (*57, 58*). Additionally, Ntrk3 and EGR1 signalling is crucial for processes underlying neuronal activity ranging from neurotransmission, synaptic plasticity and LTP to higher cognitive processes including learning and memory, social stress, and reward response (*59-62*). Moreover, *FoxJ1,* a member of Forkhead/winged-helix (Fox) family of transcription factors, play crucial role in postnatal differentiation of ependymal cells and a subpopulation of astrocytes (*63*). This subpopulation of astrocytes possesses the ability to self-renew and have neurogenic potential that can be differentiated into astrocytes, oligodendrocytes, and neurons (*63*). Additionally, studies from *FoxJ1^-/-^* neurospheres and mouse models show that FOXJ1 is important for olfactory bulb neurogenesis (*64*). Together, these results indicates that there is severe dysregulation of neuronal genes associated with various processes including neurotransmission, synaptic plasticity, learning and memory, and adult neurogenesis in *Syngap1^+/-^*mice, leading to behavioural and cognitive deficits. We propose that CSP-TTK21 treatment alleviated these behavioural and cognitive deficits in *Syngap1^+/-^*mice through restoration of epigenetic landscape (acetylation) and, consequently, the neuronal gene expression (Fig. 7).

**Fig. 7.**
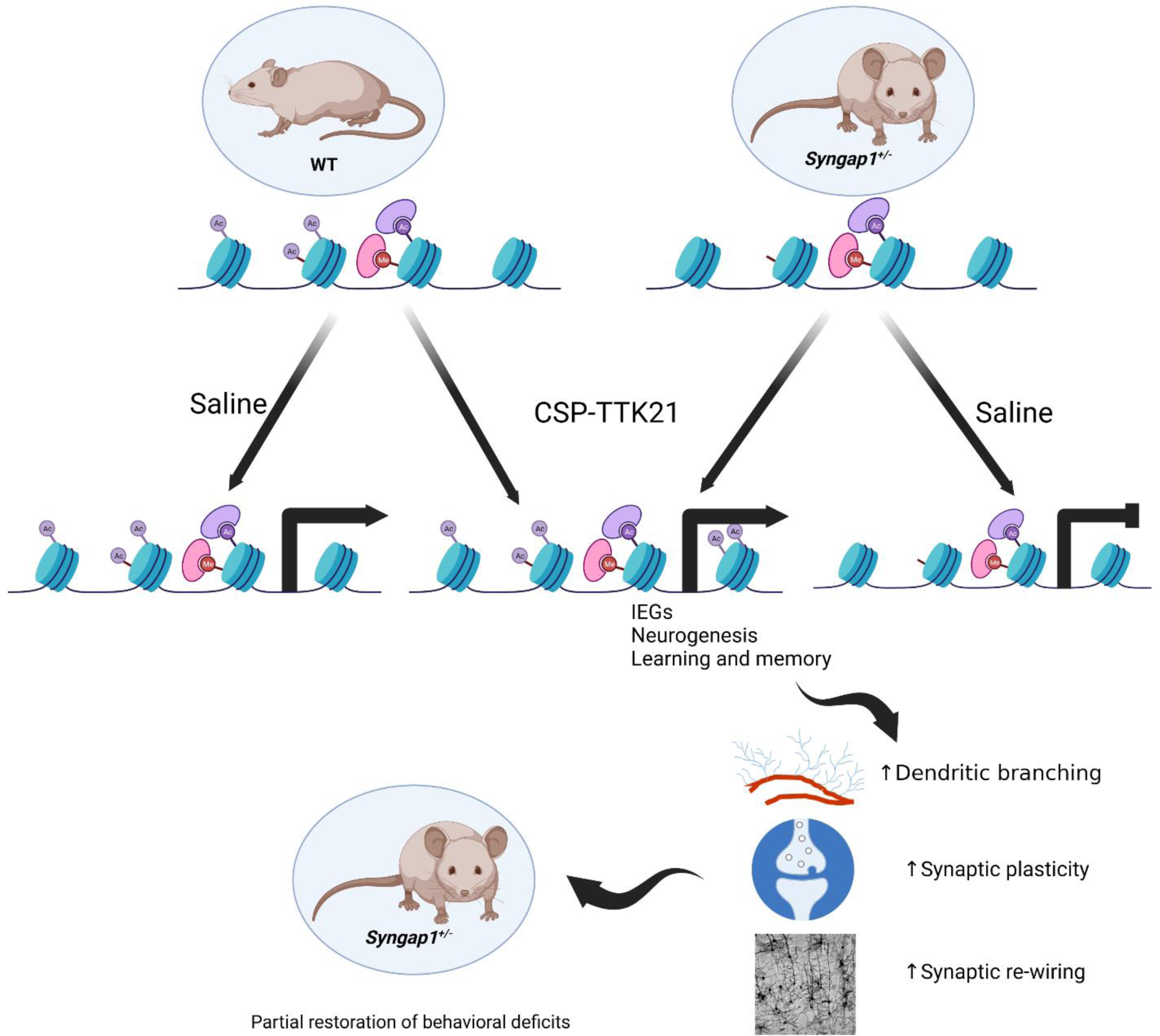
A proposed model of behavioural rescues by CSP-TTK21 in *Syngap1^+/-^* mouse. We propose that p300/CBP specific acetylation marks of histones are repressed in *Syngap1^+/-^* mouse leading to severe dysregulation of neuronal genes associated neurotransmission, synaptic plasticity, learning and memory, and adult neurogenesis, resulting in behavioural and cognitive deficits. CSP-TTK21 treatment rescues behavioural deficits by restoring transcriptomic dynamics, especially by restoring the expression of immediate early genes, genes involved in hippocampal neurogenesis and synaptic plasticity primarily through restoration of p300/CBP specific histone acetylation levels.

Our study provides a new potential therapeutic option by targeting epigenetic modifications in *Syngap1* related ID/ASD that can restore the deficits to extent which will enable the patient to lead a life less dependent on others. Recent studies corroborate our findings by targeting epigenetic modification that has shown a promising effect in the rescue of synaptic and behavioural deficits in different types of neurodevelopmental disorders (*65-67*). However, a few behavioural deficits could not be reversed by the treatment of CSP-TTK21, suggesting that, besides p300-mediated acetylation, there are other epigenetic modifications, such DNA methylation may also play critical role to establish the pathogenesis. Therefore, our studies opened a new avenue to understand the ID/IDS pathology and to design new generation therapeutics. Finally, we propose that the early therapeutic intervention should remain the primary NDD treatment strategy. Treatments during development would protect the brain from *Syngap1* hard-wired circuit damage caused by disruptions to critical neurodevelopmental processes.

## Materials and methods

### Animal Handling

C57/BL6 WT and *Syngap1^+/-^*mice were obtained from The Jacksons Laboratory (https://www.jax.org/strain/008890) and bred and maintained in Jawaharlal Nehru Centre for Advanced Scientific Research’s animal house under 12-hour dark and light cycle. Food and water were provided ad libitum. All the experiments were performed in accordance with Institutional Animal Ethics Committee (IAEC), and Committee for Purpose of Control and Supervision of Experiments on Animals (CPCSEA).

### Synthesis and characterization of CSP and CSP-TTK21

C57/BL6 WT and *Syngap1^+/-^*mice were obtained from The Jacksons Laboratory (https://www.jax.org/strain/008890) and bred and maintained in Jawaharlal Nehru Centre for Advanced Scientific Research’s animal house under 12-hour dark and light cycle. Food and water were provided ad libitum. All the experiments were performed in accordance with Institutional Animal Ethics Committee (IAEC), and Committee for Purpose of Control and Supervision of Experiments on Animals (CPCSEA).

### Synthesis and characterization of CSP and CSP-TTK21

The glucose-derived carbon nanospheres (CSP) was prepared freshly by hydrothermal process: 5g glucose in 50 ml deionized water at 180°C for nearly 12 hr. The products were isolated by centrifugation at 16000rpm, purified by repeatedly washing with water and ethanol and dried for 4 hr at 80°C. From field emission scanning electron microscopy (FESEM) imaging, we observed that the final product was spheres with an average diameter of 400 nm. TTK21 Synthesis and conjugation with CSP were done as described previously (*41*). CSP-TTK21 Activity was characterized both in cells as well as in mice by western blotting and immunohistochemistry (Fig S1).

### Immunofluorescence staining for animal tissues

Mice were administered orally with CSP and CSP-TTK21 at the dose of 20 mg/kg of their body weight. After 84 hours, mice were sacrificed by cervical dislocation. The brain was isolated, washed with PBS and transferred to 4% paraformaldehyde (PFA) for fixation at RT for 24 hrs. Then, the brain was kept in 30% sucrose solution at 4°C till processed for cryo-sectioning. Freezing of the brain was performed in optimal cutting temperature (OCT-Leica) medium for 20 min at - 20°C. 20 μm thick coronal sections were prepared from the dorsal hippocampus using a Cryotome (Leica). Sections were permeabilised in 1X PBS/2% Triton X-100 for 15 min. Nonspecific labelling was blocked by 1X PBS/0.1% Triton X-100/5% foetal bovine serum (FBS) for 30 min at 37°C. Sections were then incubated overnight with the indicated antibodies (H3K14Ac, and H4K12Ac) separately. After 3 PBS washes, sections were incubated with Alexa fluor 568 goat anti-rabbit (Invitrogen Cat # A-11004) for 1 h at room temperature. After 3 washes with 1X PBS, the nuclei were stained with Hoechst (Invitrogen Cat # H1399) for 5 min. Then sections were washes thrice and then mounted on slides. Images were acquired through confocal laser scanning microscopes LSM510 META and Zeiss LSM810. Zen 3.5 (blue edition) was used for image processing and ImageJ was used for analysis.

### Lysate preparation

#### A. Brain Lysate

Mice were sacrificed by cervical dislocation and the brain was rapidly removed and thoroughly washed with chilled 1X PBS to remove the blood samples. Cortex and hippocampal regions of the brain were dissected and homogenized in RIPA lysis buffer (150mM NaCl, 50mM Tris-Cl, 5mM EDTA, 0.10% SDS, 0.25% Sodium deoxycholate, 0.10% Triton X, pH 7.4) containing protease inhibitor cocktail (Roche # 11836170001). Homogenized samples were centrifuged at 12,000rpm at 4°C for 20 min. The supernatant was collected, and protein concentration was estimated by the Bradford method using Bio-Rad’s protein assay kit. The samples were aliquoted and stored at −80°C for further usage.

#### B. Cell Lysate

10^6^ cells were seeded in each well of 6-well-plates and kept for growth at 37°C. Cells were treated approximately at 80% confluency with different doses of CSP (10 and 50 μg/ml) and CSP-TTK21 (5, 10. 50, and 100 μg/ml) for 24 hours. Cells were harvested followed by centrifugation at 2000rpm at 4°C for 5 min. The cells were washed with chilled 1X PBS followed by centrifugation. Pellet was re-suspended in 10 times volume of obtained cell pellet in 1X RIPA lysis buffer. The cell suspension was homogenized with pipette tips and the homogenate was kept on ice for 2 hours with intermittent vertexing (2-3 secs). The Lysate was cleared by centrifugation at 13,000rpm at 4°C for 15 min, the supernatant was collected and stored at −20°C till further usage. Lysate protein concentration was determined by the Bradford method using Bio-Rad’s protein assay kit.

### SDS-PAGE and Western Blotting

Protein samples were loaded and electrophoresed through SDS-PAGE gel. The separated proteins were transferred to PVDF membrane at 25 V for 25 mins and 80 V for 1.5 hours for small and large molecular weight proteins, respectively. Post transfer, blots were stained with Ponceau stain, followed by PBS wash. After destaining, the blots were blocked with 5% skim milk (Himedia, #M530) or 5% BSA (Sigma, #A7906) for Phospho-proteins for 1 hour, washed once with PBS and probed with rabbit primary antibodies against H3 (1:5000), H3K14Ac (1:4000), H4K12Ac (1:2000), SYNGAP1 (1:1000) (Thermo Fisher Scientific, #PA 1-046), Actin (1:10000) (Thermo, #GX2781), Fmrp (1:1000) (Sigma, #F4055), Erk (1:1000) (CST, #9102), p-Erk(1:1000) (CST, #9101), mouse β-tubulin (1;2000) for overnight at 4°C. Blots were washed thrice with PBST (PBS+ Tween 20) and probed with HRP conjugated secondary antibodies, Goat anti-rabbit; 1:10000 (Abcam, #Ab97051) and Goat anti-mouse; 1:10000 (Abcam, #Ab97023) for 1 hour at RT. The blots were washed thrice with PBST and developed by Luminol chemiluminescent method using ECL western clarity solution (BioRad, #170-5061) in Syngene/Bio-Rad Gel-documentation system. Images were merged and bands quantified using ImageLab version 5.2.1 and ImageJ software, respectively.

### RNA Isolation and qPCR

Mice were sacrificed by cervical dislocation; the dorsal hippocampus was dissected out and chopped finely with a razor blade. All reagents and instruments used for RNA isolation were DEPC treated. Samples were homogenized in TRIzol reagent (Invitrogen, 15596026). RNA was isolated by the traditional phenol: chloroform method. After phenol-chloroform treatment, the aqueous phase was collected, and ethanol precipitated using absolute ethanol. The pellets were washed twice with 70% ethanol and air-dried. RNA pellets were resuspended in RNase-free water. DNase treatment was done for 30 min at 37 °C. RNA samples were further purified by chloroform/phenol extraction and ethanol precipitated. Pellets were washed twice with 70% ethanol and resuspended in RNase-free water at 65 °C for 10 mins. RNA concentration was determined by Nano-drop. 2μg of total RNA was utilized for cDNA synthesis using the MMLV-reverse transcriptase (Sigma: M 1302) and oligo-dT (Sigma: O 4387) as per the manufacturer’s instructions. qRT-PCR was carried out with TB Green Premix Ex Taq II (Takara: RR82LR) using gene-specific primers, designed, and obtained from Sigma Aldrich, India (Table 1) on a CFX96 Real-Time PCR Detection System with a C1000 Thermal Cycler (Bio-Rad). The Bio-Rad CFX Maestro Software, version 4.1, was used to analyse the data. Fold changes were estimated using the formula: 2(-[Ct Test-Ct Control]). Gapdh was considered as housekeeping gene wherever it applied.

**Table 1.**
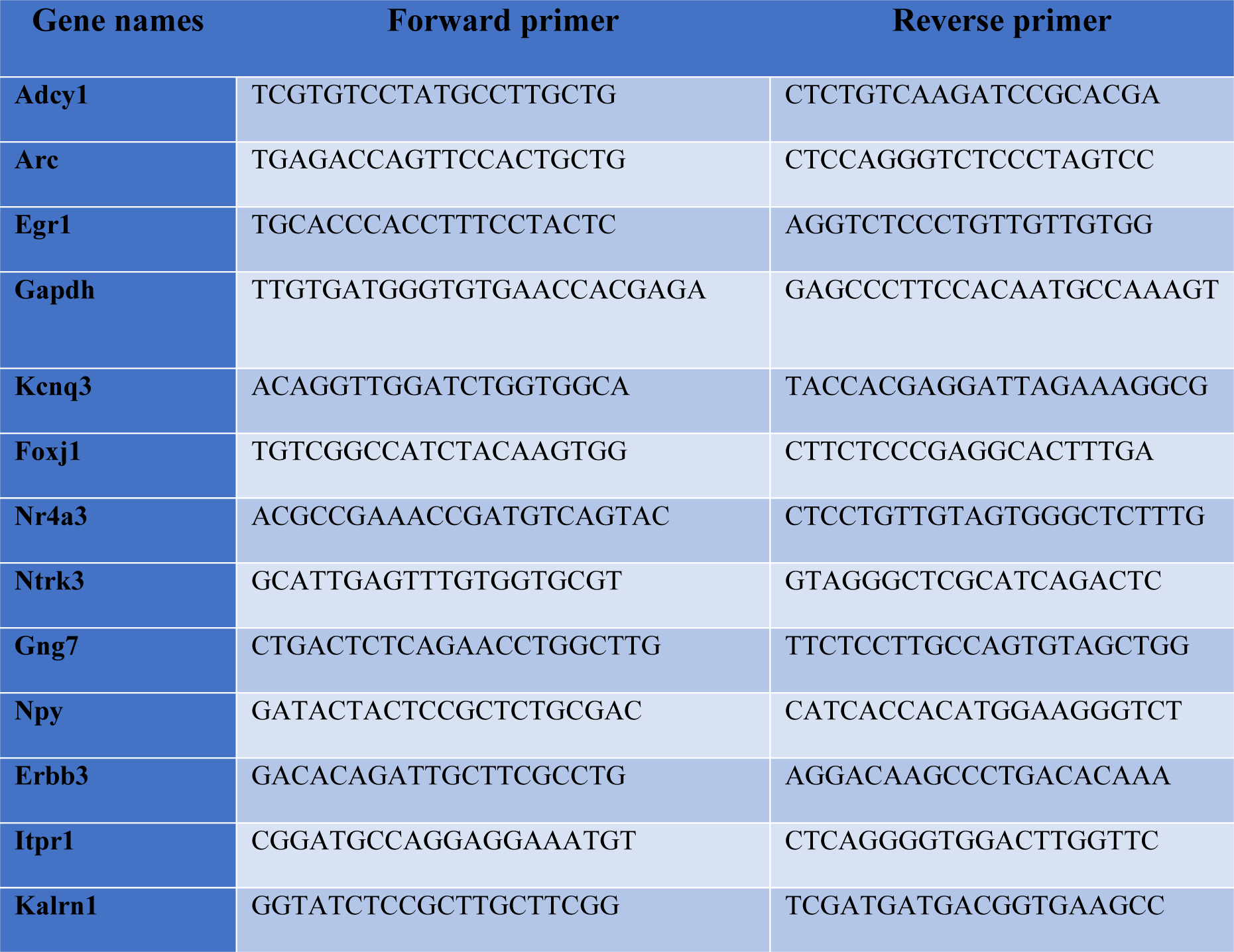
Primer list.

### Transcriptome analysis and molecular pathways

Agilent Bioanalyzer 2100 was used to ensure RNA integrity; only samples with clean rRNA peaks were used. The KAPA Stranded RNA-Seq Kit with RiboErase (KAPA Biosystems, Wilmington, MA) equipment was used to construct the libraries for RNA-seq. Agilent Bioanalyzer 2100 and Life Technologies Qubit3.0 Fluorometer were used to analyse the final library’s quality and quantity, respectively. The Illumina HiSeq 4000 (Illumnia Inc., San Diego, CA) was used for paired-end sequencing (150 bp). The genome of Mus musculus, was obtained from GENCODE and Bowtie2 with the default settings was used for indexing. Trim Galore (v 0.4.4) was used to remove adapters, and each raw Fastq file was tested for quality using FastQC. Using Samtools 1.3.1 and the “rmdup” option, PCR duplicates were eliminated. Thereafter, each raw file was aligned to the mm10 genome assembly using TopHat2 and the paired-end sequencing default parameters listed in (*68*). Following alignment, transcripts were quantified using Cufflinks, and a combined transcriptome annotation was produced using Cuffmerge. Ultimately, Cuffdiff was used to identify genes that were differentially expressed (DE). The DE gene threshold was set at log2 (fold change) <1.5 for down-regulated genes with p-value <0.05 and log2 (fold change) >1.5 for up-regulated genes. Database for Annotation Visualization and Integrated Discovery (DAVID) was used for gene ontology (GO) analysis (*69, 70*). The collection of differently expressed genes in DAVID was used for the significant enrichment test, and the Bonferroni correction method was used to select the significantly enriched biological processes. Cluster 3.0 (*71*) was used for unsupervised hierarchical clustering, with Pearson correlation and average linkage rule. Log2 transformation was applied to the gene expression data (FPKM) of all the samples. Low expression (FPKM 0.05) and invariant genes were eliminated. The genes were then centred, and clustering was performed based on gene differential expression patterns and fold change. The Java TreeView 3.0 programme was used to generate the heatmaps. GO and Pathway enrichment were performed on statistically significant differentially expressed transcripts using the DAVID programme (*69*). For additional downstream analysis, only GO terms and pathways with an FDR score of <=0.05 were taken into consideration. The selected GO terms and pathways were fed into the Pathreg algorithm (available from Theomics International Pvt Ltd, Bangalore, India) to build gene regulatory networks. To identify important nodes and edges that would be indicative of the changes in gene regulation following therapy, Cytoscape v2.8.2 was supplied with the output of the Pathreg algorithm (nodes and edges). Thereafter, these differentially expressed genes were fed into VENNY2.0 (*72*) to generate a vein diagram showing how CSP-TTK21 was able to restore the transcriptional abnormalities.

### Preparation of hippocampal slices

From WT and *Syngap1^-/+^* male and female adult mice (>6 weeks), acute brain slices were prepared as reported previously (*45*). Briefly, mice were killed by cervical dislocation, and the brain was removed and stored in a chilled sucrose-based cutting solution (124 mM Sucrose, 3 mM KCl, 24 mM NaHCO3, 2 mM CaCl2, 1.25 mM NaH2PO4, 1 mM MgSO4, and 10 mM D-Glucose). 350 µm thick horizontal hippocampal slices were cut out of the brain using an ice-cold sucrose cutting solution while it was mounted on a vibratome holder. Slices were kept in a slice chamber with artificial cerebrospinal fluid (aCSF: 124 mM NaCl, 3 mM KCl, 24 mM NaHCO3, 2 mM CaCl2, 1.25 mM NaH2PO4, 1 mM MgSO4, and 10 mM D-Glucose) for an hour in a water bath at 37°C. The slices were stored at room temperature after recovery and until usage. Following dissection, each step was carried out under continuous carbogen bubbling (5% CO2 and 95% O2; Chemix, India).

### Extracellular field recordings

Slices were moved one at a time into a submersion recording chamber that was constantly gassed with aCSF (5% CO2 and 95% O2) and kept at 34°C using in-line solution warmers (SF-28). The Schaffer-Collateral commissural pathway of the hippocampus was used to record extracellular field excitatory post synaptic potentials (fEPSPs). fEPSPs were recorded from the CA1 region of the stratum radiatum using low-resistance (3-5 M) glass electrodes filled with CSF (pH 7.3) and pulled from borosilicate glass capillaries (ID: 0.6mm, OD: 1.2mm, Harvard Apparatus) using a horizontal micropipette puller (Flaming-Brown P-97, Sutter Instruments Co., USA). A 0.1 Hz stimulation frequency was chosen. The stimulation range was set to 20–40 µs, and input–output curves were produced by varying the stimulus intensity by 20 µA per sweep in steps of 0-160 µA. A series of paired pulses separated by intervals of quarter log units, with the intervals ranging from 10 to 1000, were used to measure the paired-pulse facilitation (PPF). Recordings were discontinued and slices were discarded if the baseline activity was not stable for 15-20 minutes after the baseline activity was obtained. One train of 100Hz stimuli was used to produce long-term potentiation (LTP), and the resulting activity was monitored for 55–60 minutes. Signals were amplified using Molecular Devices’ Multiclamp 700B, filtered at 2 kHz, digitalized at 20–50 KHz using the company’s Digidata 1440A, and then saved for off-line analysis. Data was collected using pClamp 10.2 software (Molecular Devices) and analysed using Clampex 10 software.

### Behavioral paradigms

#### Handling

Mice were grouped and treated for 3 weeks (1 dosage/week) with Saline, CSP, or CSP-TTK21 prior behavioral assays. Mice were used one week after the final treatment. Mice were handled for 1-2 minutes every day for 3-5 days and proceeded for the behavior experiments. For all the behavioral experiments, mice were carried to the behavior room half an hour before the experiment. After the experiment, each mouse was returned to their home cage and the chamber was cleaned with 70 % alcohol and dried between tests.

#### Testing

##### Open Field

Each mouse was placed into one corner of an acrylic open field arena of dimensions (50 cm × 50 cm × 45), illuminated at 80-100lx. The Mouse was allowed to explore the apparatus and their activity was immediately recorded with a CCTV camera for 10 minutes. Total distance travelled, distance in the centre (20×20), number of entries in the centre, and time spent in the centre were analysed using Smart V 3.0 Software.

##### Novel Object Recognition (NOR)

Next day of OFT, each mouse was placed into one corner of the same open field arena and allowed to familiarize to two similar objects placed at two opposite corners of the arena. Familiarization was done for 2 successive days for 10 min each. After 24hr of last familiarization session, the test was performed. Testing was done for 10 min under the same condition, but with one of the familiar objects replaced with a novel object. All recordings were done with CCTV camera fitted at top of the apparatus. All the videos were analysed manually and interaction time (active contacts and mice facing the object within 2cm periphery) with each of the objects were measured. Recognition index (ratio of time spent with the novel object to total exploration time) and Differentiation index (ratio of time spent with the novel object minus the similar object to total exploration time) was calculated and plotted.

##### Social Isolation

A three-chamber rectangular transparent acrylic box (19X45 cm) was used. Central chamber had rectangular openings, allowing entrance to side chambers. Each test mouse was placed in the central chamber for 5 min with both right and left side entrance closed with heavy-weight cardboard sheets to acclimate to the apparatus. For social preference, a C57BL/6J male mouse (Stranger 1), that had no prior exposure to the test mouse, was placed inside a wire cage in one of the side chambers and an empty wire cage in the other side chamber. The wire cage was small, cylindrical, which allowed direct contact between the bars, but prevented fighting like situations. The time spent and the number of body contacts made by the test mouse with the stranger1 mouse, and the empty wire cage was analysed. For social preference, a second novel C57BL/6J male mouse (stranger 2) was introduced in the empty wire cage, the test mouse was allowed to explore both the mice (stranger1 and stranger2) for another 10 min. The time spent and the number of direct body contacts of the test mice with stranger 1 and 2 mice were analysed. Recordings were done with Sony handycam and analysed manually.

##### Morris Water Maze (MWZ)

White coloured water tank of 94 cm in diameter and 50 cm in depth was used for the test. The tank was filled with water made opaque with non-toxic white tempera paint up to a depth of 38cm. Water temperature was maintained at 25±1°C by addition of warm water when required. An escape platform 25cm^2^ in size was kept submerged just 1cm beneath the water surface in the centre of one of the quadrants (target quadrant). The platform location was kept the same during the entire training session and was removed during probe testing. Several distal extra-maze cues (a low lux bulb, a door, an empty wall, a standing experimenter) were placed around the tank and their positions remained the same during the entire training and testing sessions. A trial started by placing the mouse in one quadrant and allowed to locate the platform for 60s, once the mouse reached the platform it was left for 15s to adjust to the distal extra-maze cues. In case, the mouse was unable to locate the platform within 60s, it was manually guided to the platform by the experimenter and allowed to spend 15s to adjust to the extra-maze cues. Four trials were performed per day, 1^st^ and 4^th^ trials were started from the opposite quadrant and 2^nd^ and 3^rd^ trials were done from the right and left quadrant (from the target quadrant) respectively. After the trials, mouse was removed and dried suing a tissue paper followed by terrycloth towel and finally transferred to an empty cage kept under a lamp for at least 10min to prevent hypothermia. Each mouse was visually inspected for dryness and other issues before transferring to the home cage. All the sessions were performed at roughly the same time each day to reduce variability due to time of the day. The trials were recorded with the help of smart digital tracking system (Panlab) and were analysed using smart v.3 software. For working memory, each mouse was given 2 trials per day for 2 consecutive days, each day location of the escape platform was changed from one quadrant to other (clockwise) but was kept same for the 2 trials performed within the day. Each day the first trial was used as a training session for the new position of the escape platform, and the 2^nd^ trial was used as a test to visualize working memory. The latency to reach the target platform during the testing trials on both days were averaged out and plotted.

### DCX staining for dendritic branching

For post-treatment, adult mice were treated one dose per week for 3 weeks and after 84 hours of last treatment mice were sacrificed and brain samples were collected. For post-behaviour, mice were sacrificed, and brains were collected after completion of all the behavioural tests. The brain samples were processed for immunohistochemistry as described above. During staining, sections were incubated with DCX primary antibody at 4°C for 48 hours. After 48 hours of incubation, sections were washed thrice each time for 10 min with 1X PBS. Then sections were incubated with anti-rabbit Alexa-fluor secondary antibody for 2 hours, Hoechst was added during last 10 min of incubation. Sections were washed thrice with 1X PBS and mounting was done using anti-fade mounting media. Images were acquired with Zeiss LSM 880 confocal microscope and processed with ImageJ and NeurphologyJ software as described previously for dendritic analysis (*73*).

### Whisker-trimming and Golgi-staining

Whisker-trimming was carried out in WT and *Syngap1^+/-^* pups starting at postnatal day 7 (PND7) and was stopped at PND21 as reported previously in [11]. 18-25 mg/kg of CSP-TTK21 was administered orally post whisker trimming at PND30, PND36, and PND42. The animals were sacrificed at PND51 and transferred into Golgi-Cox staining solution (Fig 4.A). Briefly, Clean glass slides were coated with 3% Gelatin. Golgi-cox staining procedure, solution preparation, and development were carried out as described in (*74*). Animals were sacrificed at PND51 followed by dissection of the brain, washing with double distilled water (dd water), and were cut into two equal halves for better impregnation. Each hemisphere was transferred into small individual glass bottles containing Golgi-Cox solution and was kept in the dark at RT. After 24 h, each of the brain samples was transferred to a new glass bottle containing Golgi-Cox solution by pouring the Golgi-Cox solution containing the brain samples onto histological cassettes or using plastic forceps. For 7 days, the small bottles were incubated at RT in the dark. Next, the brain samples were washed with dd water and transferred into a glass bottle containing tissue-protectant solution and were kept in dark at 4 ⁰C. Post 24 h, the brain samples were now transferred into a fresh tissue-protectant solution and were stored in the dark at 4 ⁰C for 4–7 days. Following this, 80 μm thick sections were obtained in a cryostat [Leica, #CM3050 S] maintained at an object temperature (OT): −20 to −22 ⁰C and Chamber Temperature (CT): −22 to - 24 ⁰C and the sections were placed onto the gelatin-coated slides. The Golgi stained sections were developed and mounted with D.P.X mountant [Fisher Scientific, #18404] as described in (*74*).

To determine the spine and filopodia density, 8-14 dendritic segments (20–50 μm in length) per mouse were collected and considered for analysis. Pictures were visualized and elaborated with Neurolucida 360 (MBF Bioscience, Williston, VT) neuroimaging software. Selection criteria were established as reported previously to identify dendritic branches adequate for examining spine density. count/classify dendritic spines (*75*). Only transverse segments in the layers were considered for analysis. Measurements from each slice were pooled in order to obtain spine density. All measurements were performed by an experimenter blind to the experimental conditions. To determine the spine width, dendritic branches were traced using Neurolucida 360 software (MicroBrightField). Diameters of individual spines were measured by using Neurolucida explorer (MicroBrightField).

### Statistical analysis and significance test

All the statistics were carried out in Microsoft Excel 2013 and GraphPad Prism 7.02. All the graphs were made in GraphPad Prism 7.02. Data are presented as the mean ± standard error of the mean (S.E.M.) and n refers to the number of times an experiment was repeated, and N corresponds to the total number of animals used. Paired or unpaired student’s *t*-test, One-way and Two-way ANOVA with repeated measures were used to determine the significance.

## ACKNOWLEDGEMENTS

This work was supported by the Jawaharlal Nehru Center for Advanced Scientific Research (JNCASR), J C Bose Fellowship, Department of Science and Technology (DST), India (grant no. SR/S2/JCB-28/2010) and SERB (grant no. CRG/2021/000356) to TKK, AKS is supported by the JNCASR Research Fellowship.

## Author contributions

Conceptualization: T.K.K. and J.P.C. Methodology: A.K.S., I.J., N.M.N.R., P.S., E.M., M.V., and S.B. Investigation: A.K.S., I.J., N.M.N.R., and S.P. Visualization: A.K.S., I.J., N.M.N.R., S.P., and M.V. Funding acquisition: T.K.K. and J.P.C. Supervision: T.K.K., S.B., and J.P.C. Writing–original draft: A.K.S. and T.K.K. Writing–review and editing: A.K.S., S.B., J.P.C and T.K.K.

