## Supplemental File for "Epigenetic Modulation to perturb the *SYNGAP1* Intellectual Disability (ID) that ameliorates synaptic and behavioural deficits"

### Supplementary Materials.

Fig. S1 to S8

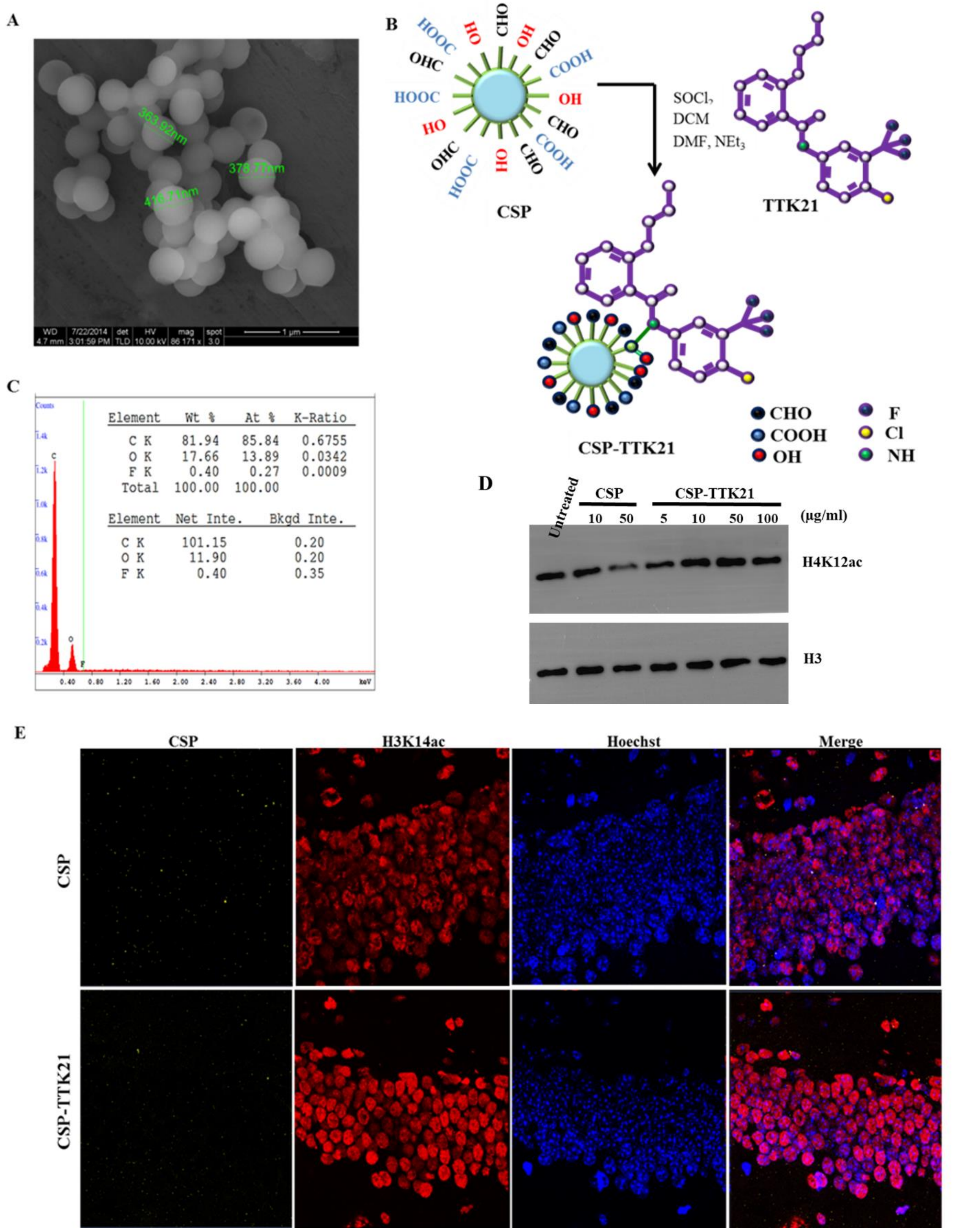


**Fig. S1. Synthesis and characterization of CSP-TTK21.** (A) FESEM image of CSP, (B) Schematic showing the synthesis of CSP-TTK21, (C) the presence of fluorine upon EDX analysis of CSP-TTK21 confirming the conjugation of TTK21 with CSP. (D) Increased H3K14 acetylation upon immuno-blotting of CSP and CSP-TTK21 treated SHSY-5Y cells as well as (E) Induction of H3K14 acetylation in dorsal hippocampus of mouse brain treated with CSP-TTK21, shows CSP-TTK21 is functionally active. Scale (A) 1µm and (E) 10µm.

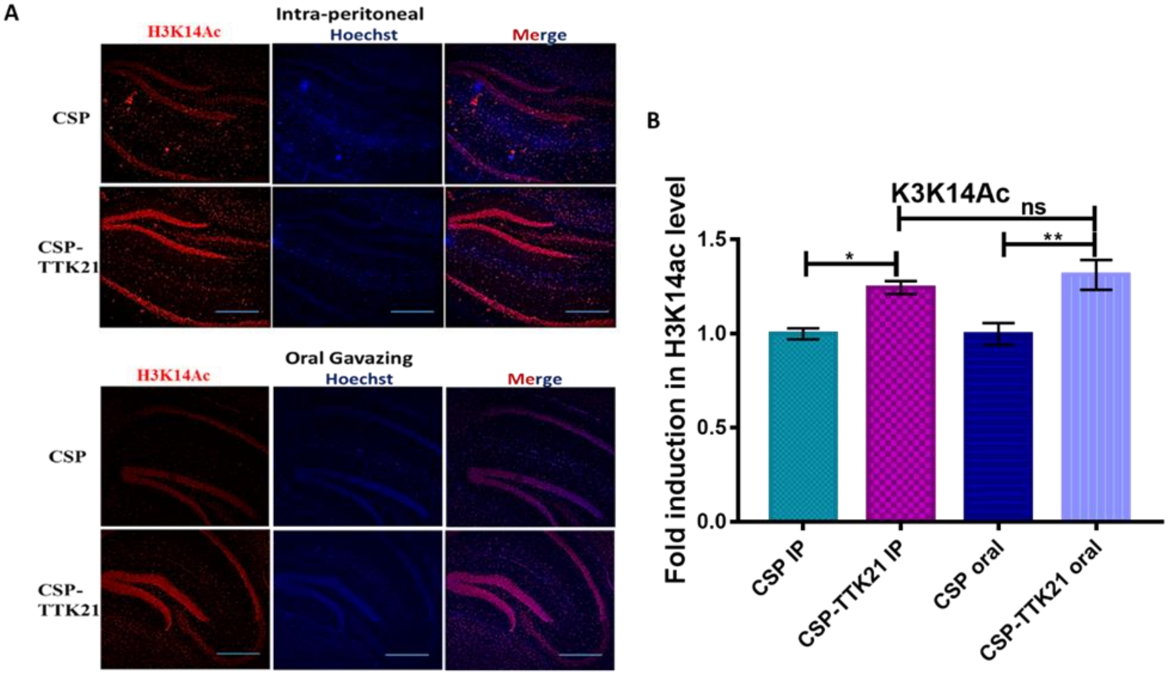


**Fig. S2. CSP-TTK21 induces histone acetylation upon oral gavazing. (A)** In both cases IP and Oral administration, acetylation was increased in the hippocampal region as compared to CSP-control (n=4). Scale bar 200µm. (B) Quantitation of fold induction in histone acetylation level upon Oral and IP administration of CSP-TTK21 (normalized to CSP). Error bars represent standard error of mean (SEM). Student's t-test was done for significance test, $*p<0.05$ and $**p<0.01$.

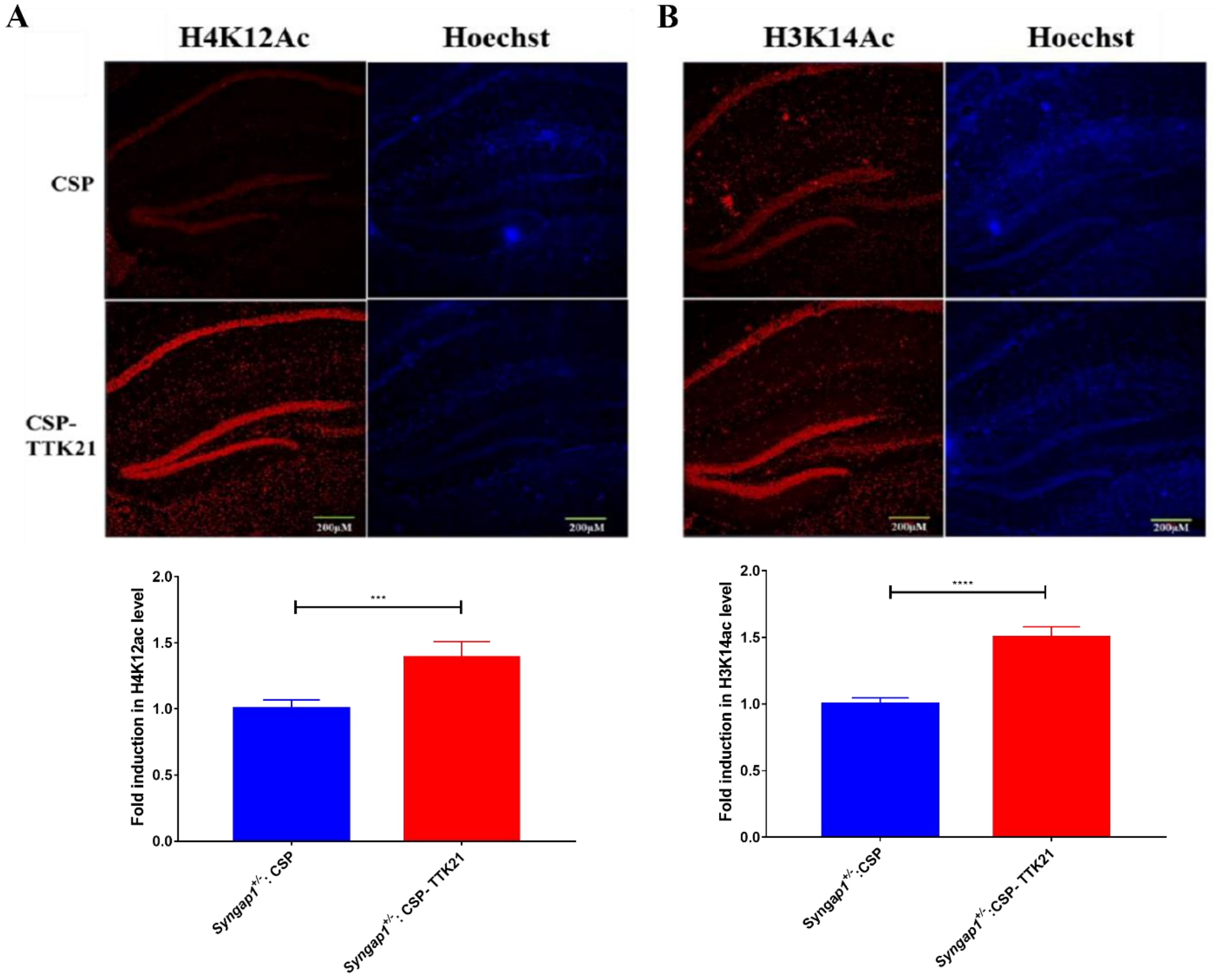


**Fig. S3. CSP-TTK21 treatment increased histone acetylation in *Syngap1*$^{+/-}$ mouse.** Representative confocal images showing induction of (A) H4K12Ac levels and (B) H3K14Ac levels in dorsal hippocampus of *Syngap1*$^{+/-}$ mice upon treatment with CSP-TTK21 as compared to CSP vehicle control. Error bars represent standard error of mean (SEM). Student's t-test was done for significance test, ***p<0.001, and ****p<0.0001.

**A**

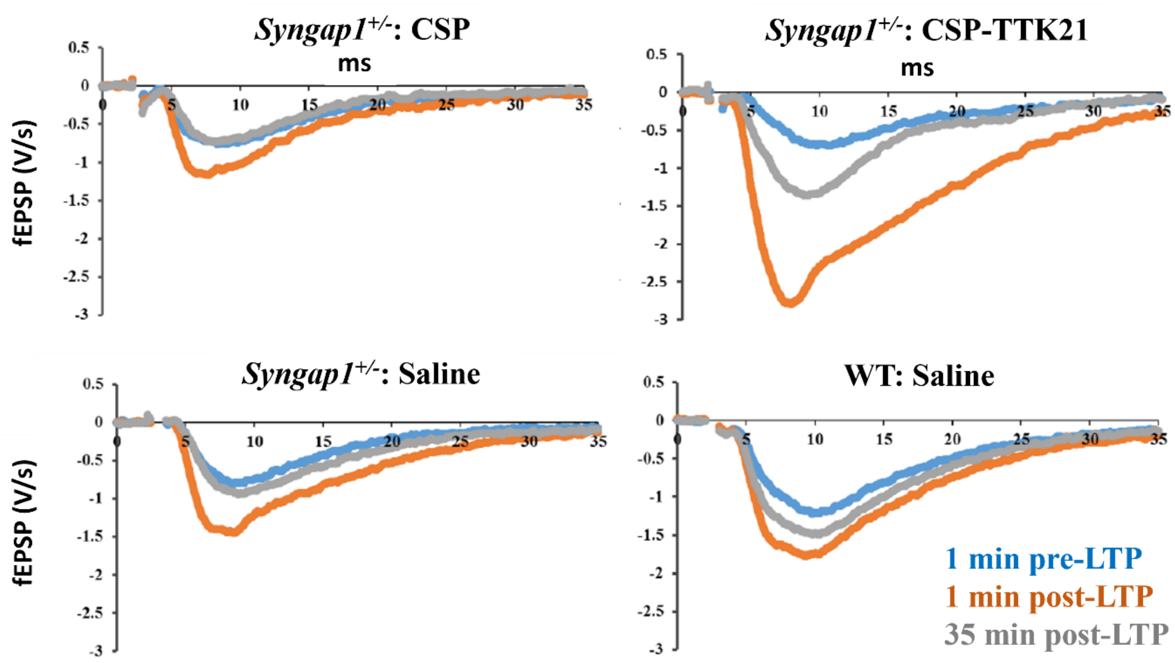


**B**

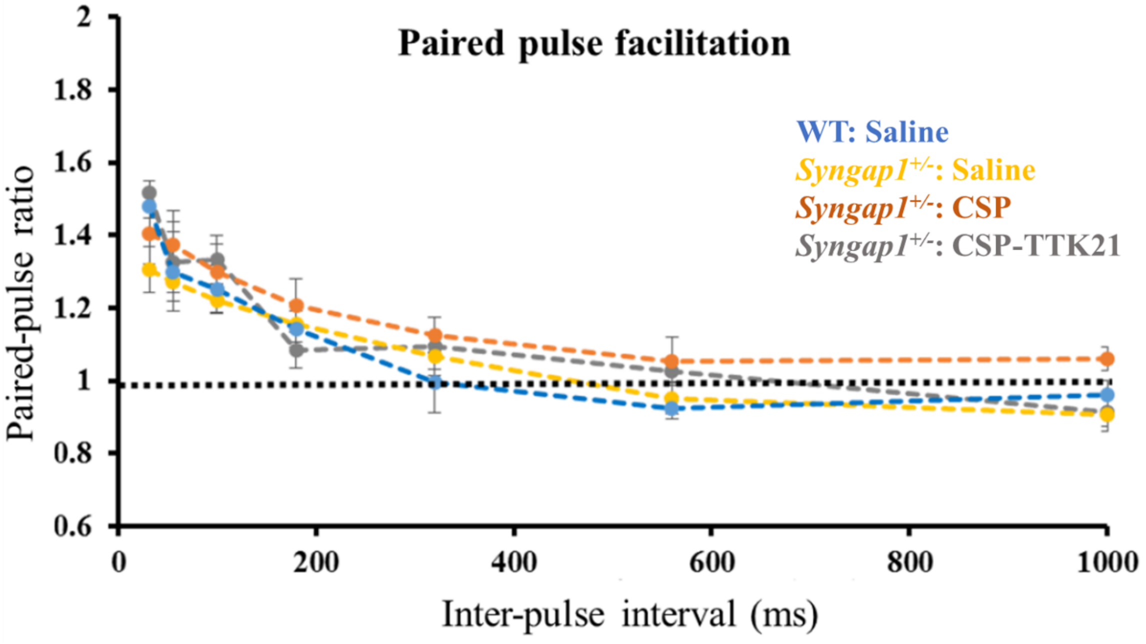


**Fig. S4. CSP-TTK21 induces LTP potentiation, not paired pulse response (PPR).** (A) Representative sample traces for each group, taken 1 min before (blue), 1 min after (orange) and 35 min after (grey) LTP induction. (B) Two presynaptic spikes were evoked simultaneously at different time interval and the ratio of the two post-synaptic response (fEPSP2/fEPSP1) were measured and plotted against time interval. No significant alteration was seen in paired pulse facilitation, $p>0.06$. Significance was tested by Two-way ANOVA (repeated measures). Error bars represent SEM. n= 4 slices.

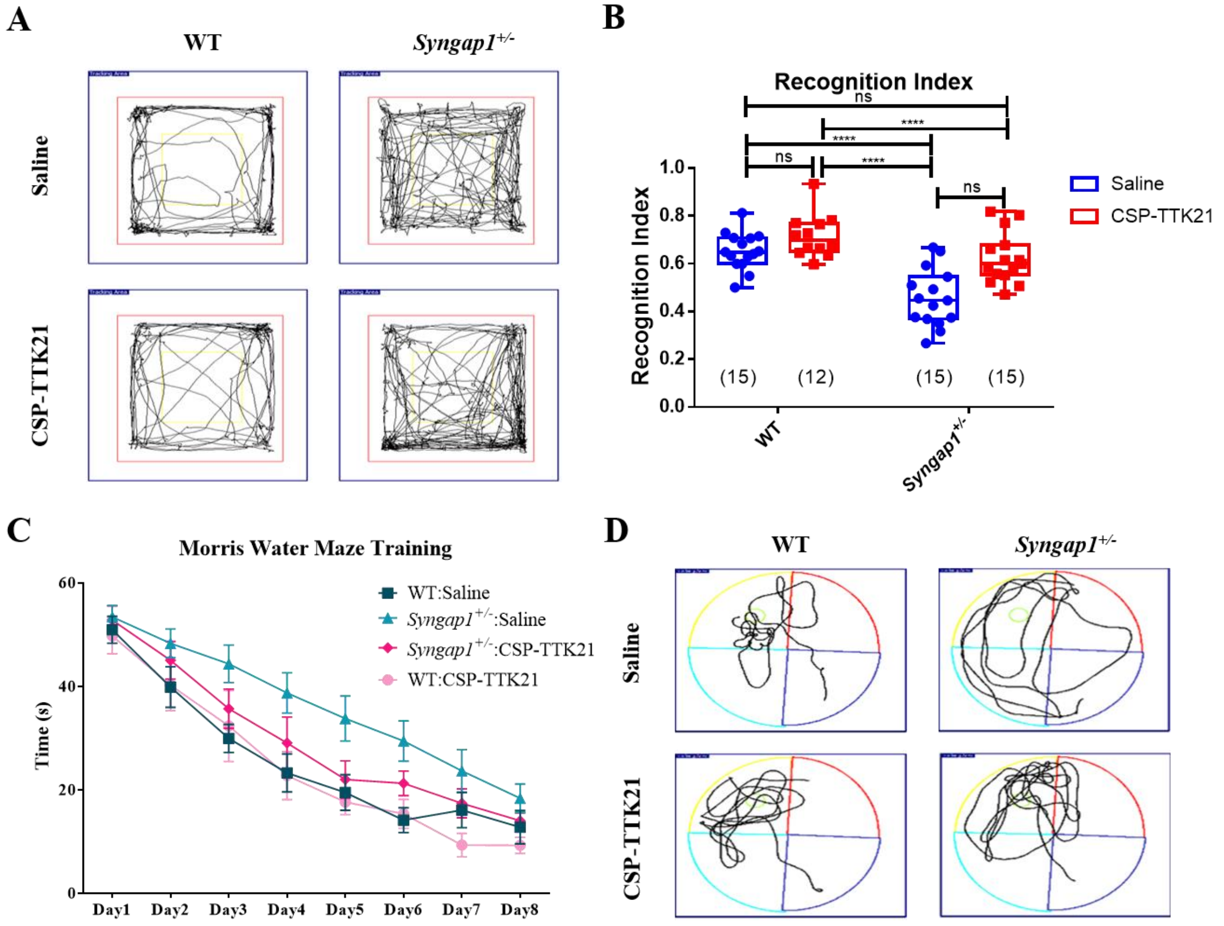


**Fig. S5. CSP-TTK21 effects on behavioural measures in *Syngap1*$^{+/-}$ mouse.** (A) OFT representative tracks. (B) Recognition index (RI) in Novel object recognition memory, reduced RI in *Syngap1*$^{+/-}$ mice representing impaired object recognition memory. CSP-TTK21 treatment increases RI in *Syngap1*$^{+/-}$ mice. (C) The latency time across different days of MWM training (12-15 mice/group). (B) Representative traces from each group during probe test depicting localized movement in the target zone (yellow) in CSP-TTK21 treated *Syngap1*$^{+/-}$ mice as compared to random movement in *Syngap1*$^{+/-}$:Saline group.

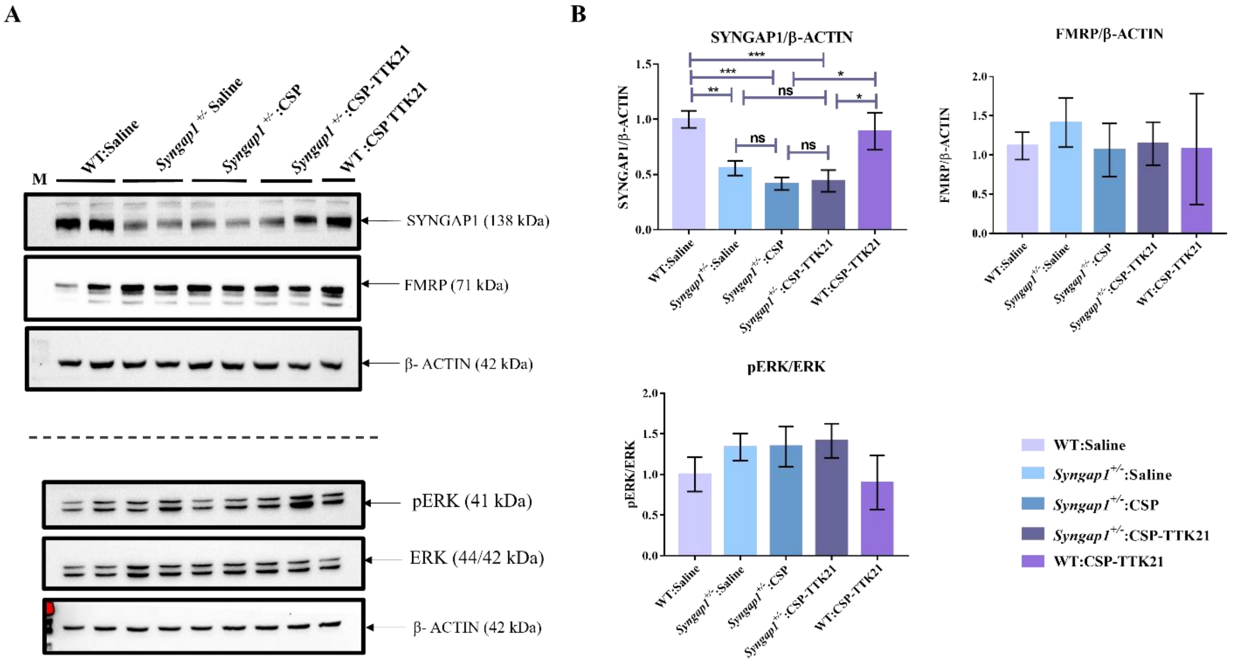


**Fig. S6. Western blotting from mice lysate after treatment with CSP or CSP-TTK21**. Mice were sacrificed and tissue lysates were prepared after 3 days of CSPTTK21 treatment (4-6 mice/group). (A) Representative blot images and (B) Quantifications showing no effect of CSP-TTK21 on SYNGAP1, p-ERK/ERK, and FMRP levels.

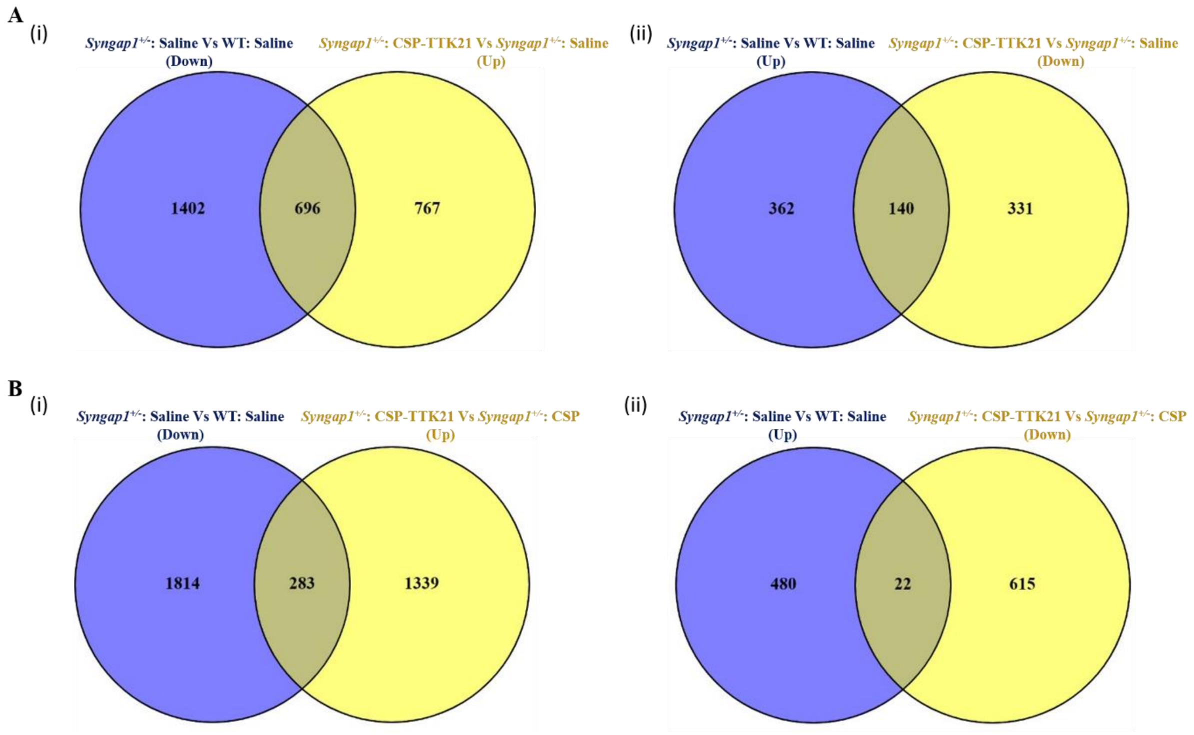


**Fig. S7. Venn diagram showing effect of CSP-TTK21 in restoration of significantly deregulated genes across different treatment conditions**.

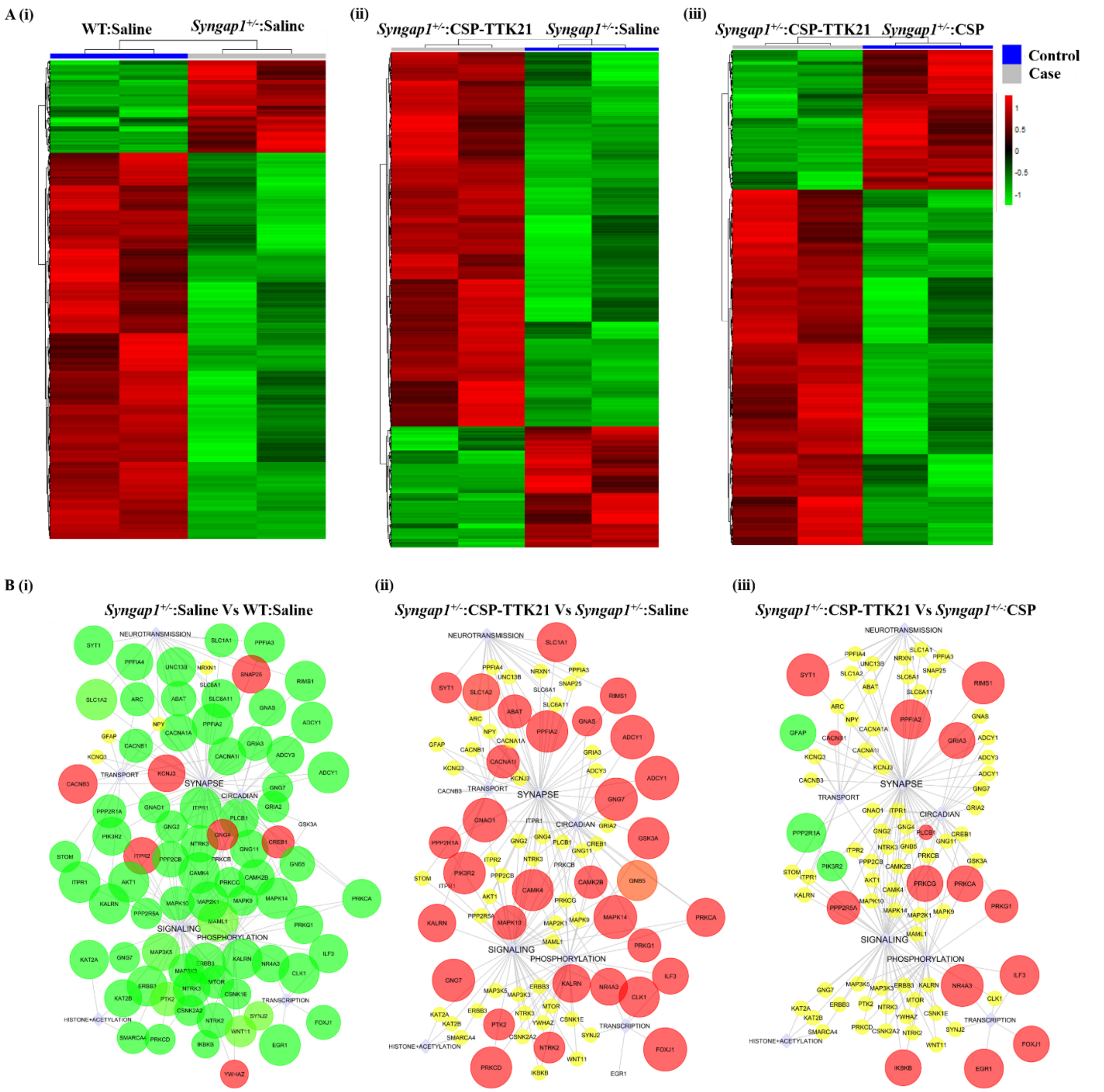


**Fig. S8. CSP-TTK21 restores gene expression and signaling pathways that were altered in *Syngap1*$^{+/-}$ mouse.** (A) Heatmaps representing the differentially expressed transcripts across various conditions. (i) *Syngap1*$^{+/-}$: Saline Vs WT: Saline, (ii) *Syngap1*$^{+/-}$: CSP-TTK21 Vs *Syngap1*$^{+/-}$: Saline, and (iii) *Syngap1*$^{+/-}$: CSP-TTK21 Vs *Syngap1*$^{+/-}$: CSP. Blue colour represents the control group and grey represents the case group (effect) (A) Affected pathways in *Syngap1*$^{+/-}$ mouse (*Syngap1*$^{+/-}$: Saline Vs WT: Saline). (B) Effect of CSP-TTK21 treatment (*Syngap1*$^{+/-}$: CSP-TTK21 Vs *Syngap1*$^{+/-}$: Saline). (C) Direct effect of CSP-TTK21 (*Syngap1*$^{+/-}$: CSP-TTK21 Vs *Syngap1*$^{+/-}$: CSP).
